# Control of physical and biochemical parameters influencing exogeneous cargo protein association to extracellular vesicles using lipid anchors enables high loading and effective intracellular delivery

**DOI:** 10.1101/2024.08.28.610030

**Authors:** Antonin Marquant, Jade Berthelot, Claudia Bich, Zeineb Ibn Elfekih, Laurianne Simon, Baptiste Robin, Joël Chopineau, David Tianpei Wang, Samuel Jay Emerson, Aijun Wang, Clément Benedetti, Simon Langlois, Laurence Guglielmi, Pierre Martineau, Anne Aubert-Pouëssel, Marie Morille

## Abstract

Despite biomolecule delivery is a natural function of Extracellular Vesicles (EVs), low loading of exogenous macromolecules such as proteins into EVs limits their interest as convincing protein delivery systems for health applications. In this context, lipid-anchorage of exogenous cargo into EV membrane recently emerged as a promising option to enable their vectorization into cells. Nevertheless, this option was not explored for protein intracellular delivery, and further characterization of critical parameters governing the association of a lipid-anchored cargo protein to EVs stills needed to confirm the relevance of this anchorage strategy. Therefore, we sought to identify these parameters in a precise and quantitative manner, using bulk and single nanoparticle analysis methods to identify protein loading capacity and subsequent intracellular delivery. We identified incubation temperature, cargo concentration, Lipid Anchor (LA) structure (lipid nature, linker) and EV origin as critical factors influencing maximal EV loading capacity. Precise control of these parameters enabled to load cargo protein close to EV saturation without hindering cellular delivery. Structural properties of LA influenced not only cargo protein/EV association, but also intracellular delivery into different carcinoma cell lines. By thoroughly characterizing Lipid-PEG-protein anchorage, this study evidences the interest of this tunable and controllable approach for efficient EV protein delivery.

**Graphical abstract:** 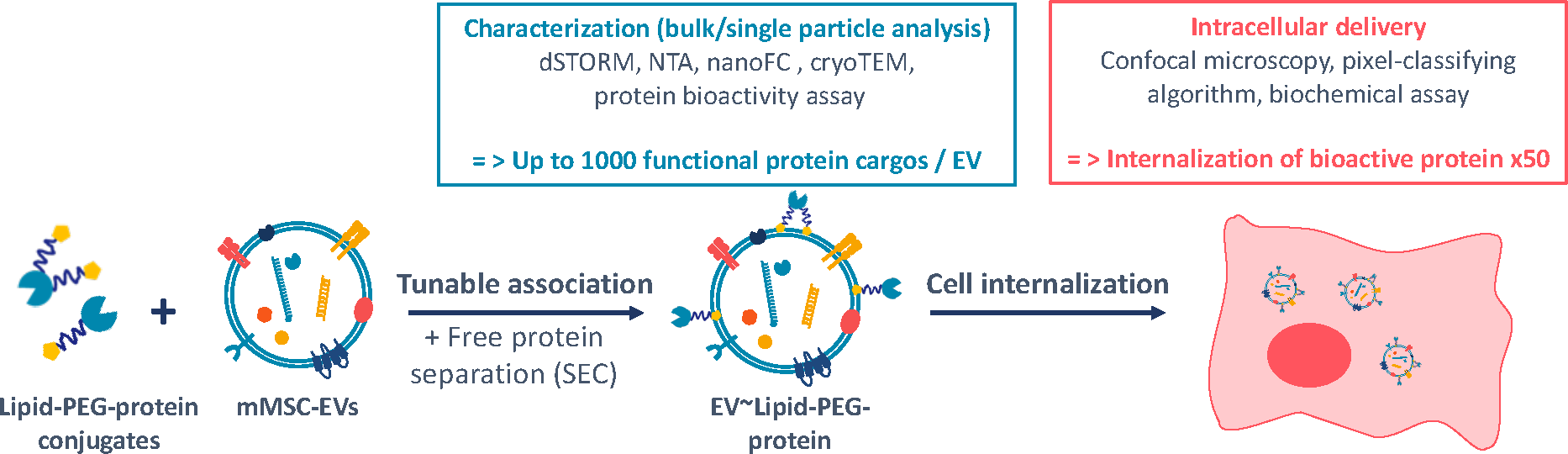

## 1. Introduction

Therapeutic proteins are powerful and fast-acting drugs that have proved their efficacy for many years in the treatment of a wide range of pathologies such as cancers^1–3^, infectious or genetic diseases (HIV infection^4^, cystic fibrosis^5^, etc.) or auto-immune pathologies^6,7^. By contrast to small molecules, proteins have pharmacological effects at very low concentration and are highly specific for their targets^8^. As a consequence, development of therapeutic proteins has soared over the two last decades leading proteins such as antibodies to be top-tier drugs^9^ and this trend is expected to continue^10^. However, despite their very interesting intrinsic properties, proteins also struggle with important limitations hindering their therapeutic potential. Delivery is one of the main challenges to overcome, as these fragile macromolecules face numerous barriers before reaching their target, especially if intracellular delivery is sought. For molecules of this size, crossing the cell membrane is impossible, and no delivery system capable of fully resolving this problem has yet been developed. As a result, most of currently marketed therapeutic proteins target extracellular sites such as membrane proteins^11^, limiting effective therapeutic applications.

Among promising options, natural nanoparticles such as Extracellular Vesicles (EVs) have been recently considered^12^. EVs are produced by the totality of known cells and organisms and have the ability to vectorize biomolecules from nucleic acids to proteins, making them a highly attractive alternative to synthetic nanoparticle-based therapies^13^. EVs have been successfully used to deliver several proteins^14–20^ relying either on endogenous (e.g., using cell engineering for protein fusion^18,21–23^) or exogenous (e.g., using isolated EV submitted to membrane permeabilization processes^20,24^) loading methods. Using endogenous methods, reported loading efficiencies are usually limited below 200 proteins/EV^21^ and engineering efficiency (i.e., the proportion of EV loaded as compared to total EV population) is generally below 30%^21,25–27^. Moreover, these strategies imply time-consuming cell genetic manipulations and restrict the loading to certain EV sub-populations (e.g., CD63 positive EV). In contrast, exogenous methods rely on physical destabilization of EV membranes. These approaches have the disadvantage of producing protein aggregates that form during this harsh process and may be confused with EV. Moreover, the volume occupied by EVs, even at high concentration (<5.10^11^EV/ml), is too low to enable an efficient loading only based on cargo diffusion into EVs. Finally, it is difficult to compare all these approaches due to the lack of characterization and comparable metrics. Thus, development and characterization of efficient protein loading approaches for EVs is urgently needed.

In this context, the addition of a Lipid Anchor (LA) to cargo is an appealing option to increase EV loading by exploiting hydrophobic interactions with the phospholipid EV membrane in mild conditions. Moreover, this strategy can be considered more versatile, especially compared to fusion with EV proteins, as all EV sub-populations are enclosed by hydrophobic membrane. This approach has been successfully used for some types of cargos such as small chemicals^28–31^ or nucleic acids^32–36^. However, cargo proteins were barely reported and never for intracellular delivery and parameters able to influence cargo protein association to EVs remain to be investigated. Some studies suggested the impact of the lipid nature of LA but only for small cargos^29,37^ and potential effects from other parameters (e.g linker) were never investigated, especially for intracellular administration of a protein.

This study intends to characterize the LA-cargo protein association to EVs by using biochemical readouts, state-of-the-art single-particles analysis methods and algorithmic pixel-classifying analysis of confocal microscopy images. We assumed that control of fundamental parameters governing this loading could turn EVs into effective intracellular protein delivery systems while maintaining both protein and EV properties. We investigated the influence of physical parameters (cargo concentration and incubation temperature), and LA structure (lipid nature and PEG linker molecular weight). All these parameters appeared crucial for the maximal loading capacity of EVs, the stability of the association or the efficiency of intracellular delivery. By controlling this process, we obtained a very high loading capacity equal to ≈1000 cargo proteins per EVs. Our results therefore evidenced this strategy as a new interesting option for protein intracellular delivery, opening promising opportunities for various therapeutic applications.

## 2. Results

### 2.1. Characterization and tunability of the association of Lipid-PEG-cargo to EVs

#### 2.1.1 Cargo protein concentration and incubation temperature are key parameters controlling maximal association to EVs and enable high loading efficiency

A commercial source of EVs (from Everzom company) produced by murine mesenchymal stem cells (mMSC) and extracted by tangential flow filtration followed by a steric exclusion chromatography step was characterized (Supplementary Figure 1). These EVs exhibited expected EV features (i.e., morphology, size and antigen markers). Horse Radish Peroxidase (HRP) was chosen as model cargo protein for its representative size (Hydrodynamic Radius ≈3nm and 45kDa) and extremely accurate enzymatic readout, enabling us to validate the maintain of protein activity during this process. Thus, Lipid-PEG-HRP (namely DSPE-PEG_1000_-HRP, DSPE-PEG_2000_-HRP, DSPE-PEG_5000_-HRP and CLS-PEG_2000_-HRP) were obtained after reaction of maleimide-functionalized HRP and Lipid-PEG-SH anchors as described in section 5.3.1 and summarized in Supplementary Figure 2B. Lipid-PEG-HRP conjugates were characterized by HPLC-MS (Supplementary Figure 2C). Conservation of enzymatic activity of HRP was controlled after conjugation (Supplementary Figure 2D). Steric Exclusion Chromatography (SEC) was used to remove all non-associated cargo protein from samples and to measure accurate association capacity (Figure 1A). Fractions 3, 4 and 5 were identified as EV-containing fractions (13±3%, 28±4%, and 16±2%, respectively) while negligeable amount of free Lipid-PEG-HRP was found in these fractions (0%, 0% and 2±1.4%., respectively). Thus, the F4 fraction was selected to ensure that no free lipid-PEG-HRP could contaminate samples and lead to an overestimation of the association calculation.

**FIGURE 1.**
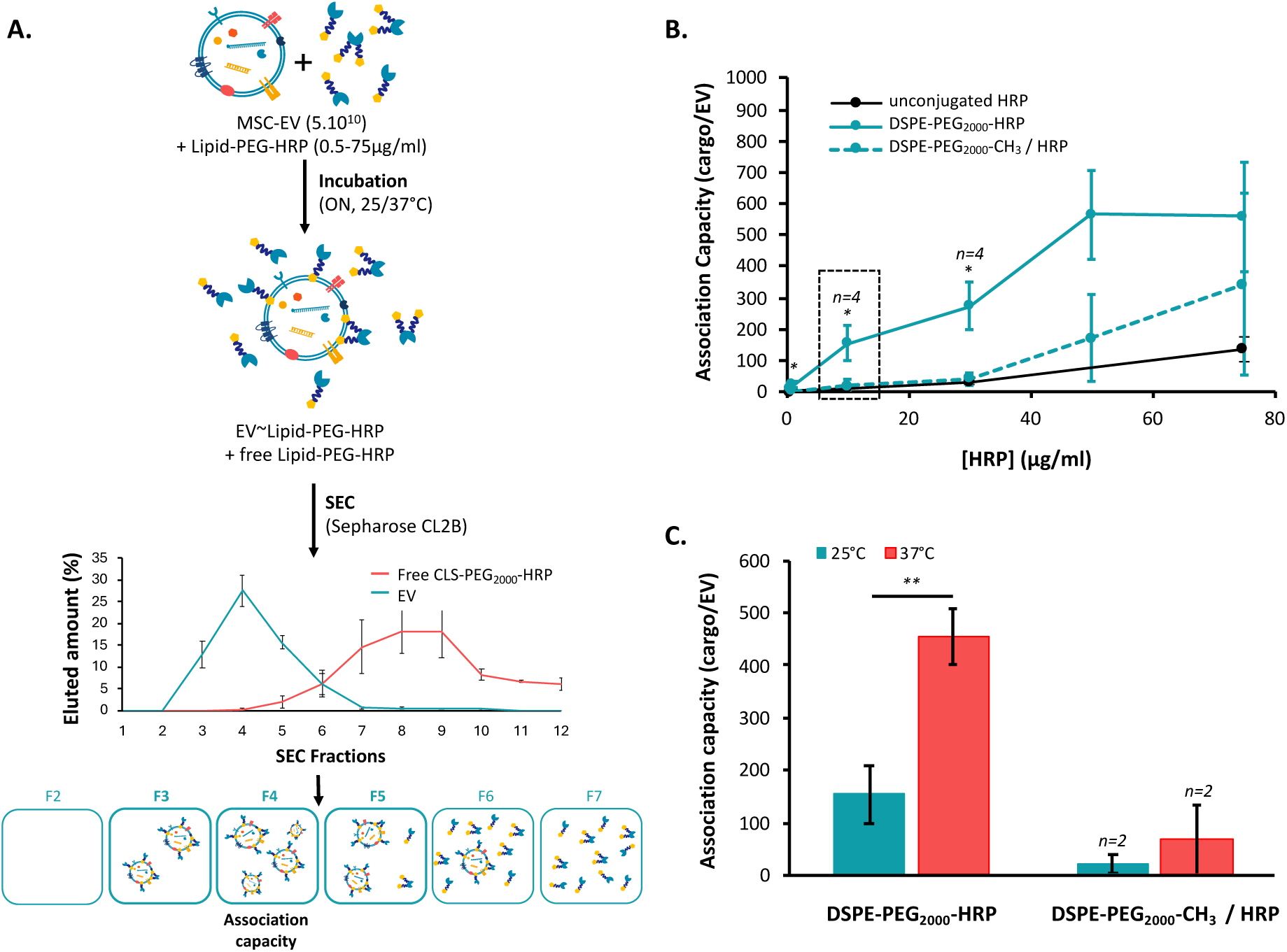
Influence of Lipid-PEG-cargo concentration and temperature on the association with EVs. **(A)** Schematic representation of EV / Lipid-PEG-HRP association workflow (association, separation), not at scale. The 4^th^ fraction of SEC was selected to calculate association capacity. **(B)** Number of biologically active Lipid-PEG-HRP per EV *vs.* HRP concentration after incubation of 5.10^10^ EV/ml with unconjugated HRP, unconjugated DSPE-PEG_2000_-CH_3_ / HRP, and DSPE-PEG_2000_-HRP at 25°C overnight. n=3 if unspecified. Significance of p-values as compared to DSPE-PEG_2000_-CH_3_ / HRP control are given above each point as well as between two successive points. **(C)** Number of biologically active HRP conjugated per EV *vs.* temperature during incubation of 5.10^10^ EV/ml with 10µg/ml of DSPE-PEG_2000_-CH_3_ / HRP or conjugated DSPE-PEG_2000_-HRP at 25°C or 37°C overnight. p-values were calculated using a two-tailed Welch test. *n.s*>0.05, *<0.05, **<0.01.

Finally, two negative controls were added in association experiments to ensure that the retention of HRP with EVs in SEC fractions was the result of their association and not an artifact due to HRP and Lipid-PEG aggregates: 1) unconjugated HRP with EVs and 2) Lipid-PEG-CH_3_ + HRP with EVs. For this last condition, the N-terminal methoxy function (CH_3_) does not allow the covalent attachment of Lipid-PEG to mal-HRP.

We first investigated the influence of HRP concentration and incubation temperature to identify optimal association conditions. First experiment set was done with a fixed EV concentration (5.10^10^ EV/ml) incubated with various concentrations of DSPE-PEG_2000_-HRP (from 0.5µg/ml to 75µg/ml) at 25°C (Figure 1B). After SEC separation, association capacity was firstly calculated based on the assumption of a homogeneous distribution of HRP among EVs in fraction F4. Association capacities of controls remained constantly below 50 HRP/EV without significant variation between 0.5µg/ml and 30µg/ml. Above this concentration, association capacities started to increase, reaching 341±289 HRP/EV and 134±39 HRP/EV at 75µg/ml, respectively. In contrast, DSPE-PEG_2000_-HRP association to EVs started from 8±1 HRP/EV at 0.5µg/ml, and increased progressively to 21±10 HRP/EV at 1µg/ml, 154±56 at 10µg/ml and 272±76 at 30µg/ml to finally reach a plateau of 564±144 HRP/EV at 50µg/ml. Number of HRP associated with EVs was significantly higher than controls when HRP was conjugated with DSPE-PEG_2000_ for each tested concentration (excepted at 75µg/ml, probably due to SEC column saturation).

Then, we evaluated the influence of incubation temperature (25°C or 37°C) at 10µg/ml of DSPE-PEG_2000_-HRP (Figure 1C), leading to a rise in association from 154±56 HRP/EV to 456±53 HRP/EV while no significant increase was observed in the control. Noticeably, this value of 456±53 HRP/EV was close to EV saturation previously observed for higher DSPE-PEG_2000_-HRP concentrations (50µg/ml) at 25°C. These results suggest that increase in EV membrane fluidity facilitates LA-cargo insertion. This incubation at 37°C was not deleterious for EVs based on mode, concentration and cryoTEM observations (Supplementary Figure 3). Therefore, association conditions were set to a Lipid-PEG-HRP concentration of 10µg/ml incubated at 37°C overnight with 5.10^10^ EV/ml for subsequent experiments.

#### 2.1.2. Lipid-PEG anchor structure influences the cargo protein association to EVs

We compared three different MW of PEG linkers for the same lipid anchor (DSPE) with 1000g/mol, 2000g/mol and 5000g/mol (Figure 2A). Similarly, cholesterol-PEG_2000_ (abbreviated in CLS-PEG_2000_) was compared to DSPE-PEG_2000_ (Figure 2B). DSPE-PEG-HRP conjugates were efficiently associated to EVs without significant differences depending on the PEG MW, with an association capacity of 373±110 HRP/EV, 255±57 HRP/EV and 329±86 HRP/EV using PEG 1000, 2000 and 5000g/mol, respectively. Surprisingly, an association of 898±247 HRP/EV was obtained for CLS-PEG_2000_-HRP, above the EV surface saturation observed with DSPE-PEG_2000_-HRP in the same conditions (10µg/ml of LA-HRP incubated at 37°C overnight).

**FIGURE 2.**
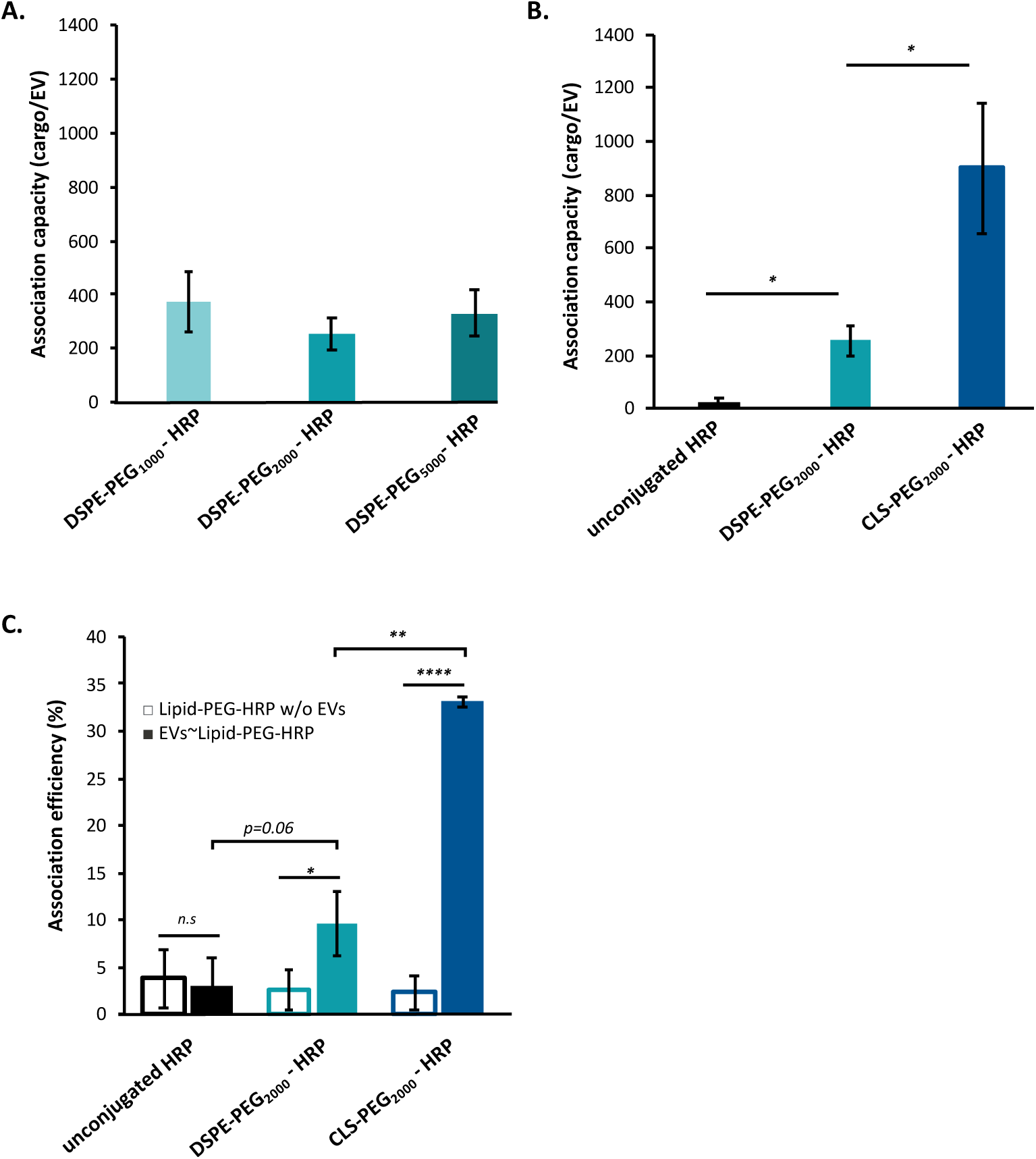
Influence of PEG linker MW, lipid anchor nature and EV type on the association between EVs and Lipid-PEG-HRP. Association efficiency according to **(A)** the PEG MW of DSPE-PEG anchor or **(B)** the lipid nature of Lipid-PEG_2000_ anchor after incubation of 5.10^10^ EV/ml with 10µg/ml of HRP incubated overnight at 37°C. 4^th^ SEC fraction was considered to calculate this association capacity (n=3). **(C.)** Loading efficiency defined as percentage of HRP Lipid-PEG-HRP retained with EVs (all EV SEC fractions were considered) after incubation of 10µg/ml of unconjugated HRP, conjugated DSPE-PEG_2000_-HRP and conjugated CLS-PEG_2000_-HRP with (n=3) 5.10^10^ EV/ml or without (n=4) EV incubated overnight at 37°C. p-values were calculated using a two-tailed Welch test. n.s>0.05, *<0.05, **<0,01, ***<0,001, ****<0,0001.

This high loading capacity conferred to this process an important association efficiency (i.e., the percentage of total available cargo that has been associated with EVs compared to the total amount introduced into the incubation media^27^). After an incubation of 10µg/ml of unconjugated HRP, DSPE-PEG_2000_-HRP or CLS-PEG_2000_-HRP with EVs, association efficiencies of 3.1±2.9%, 9.7±3.3% and 33.2±0.5% were obtained, respectively (Figure 2C). By considering that only 60% are collected after SEC, the corrected loading efficiency is superior to 50% using CLS. Therefore, we selected CLS-PEG_2000_ as reference Lipid-PEG anchor for subsequent experiments and characterization.

#### 2.1.3. Single particle analysis unveils EV∼CLS-PEG_2000_-cargo association heterogeneity

We fluorescently labeled HRP for single particle analysis measurements and co-localization observations in order to have an overview of the repartition of CLS-PEG_2000_-cargo among EVs. Unconjugated HRP and CLS-PEG_2000_-HRP were conjugated with Alexa-Fluor 488 TFP ester to obtain unconjugated HRP-A488 and CLS-PEG_2000_-HRP-A488. NTA analysis using Zetaview-Twin showed no fluorescent particles for EVs incubated with unconjugated HRP-A488 while 59±14% of detected particles were fluorescent in EV∼CLS-PEG_2000_-HRP-A488 samples (Figure 3A).

**FIGURE 3.**
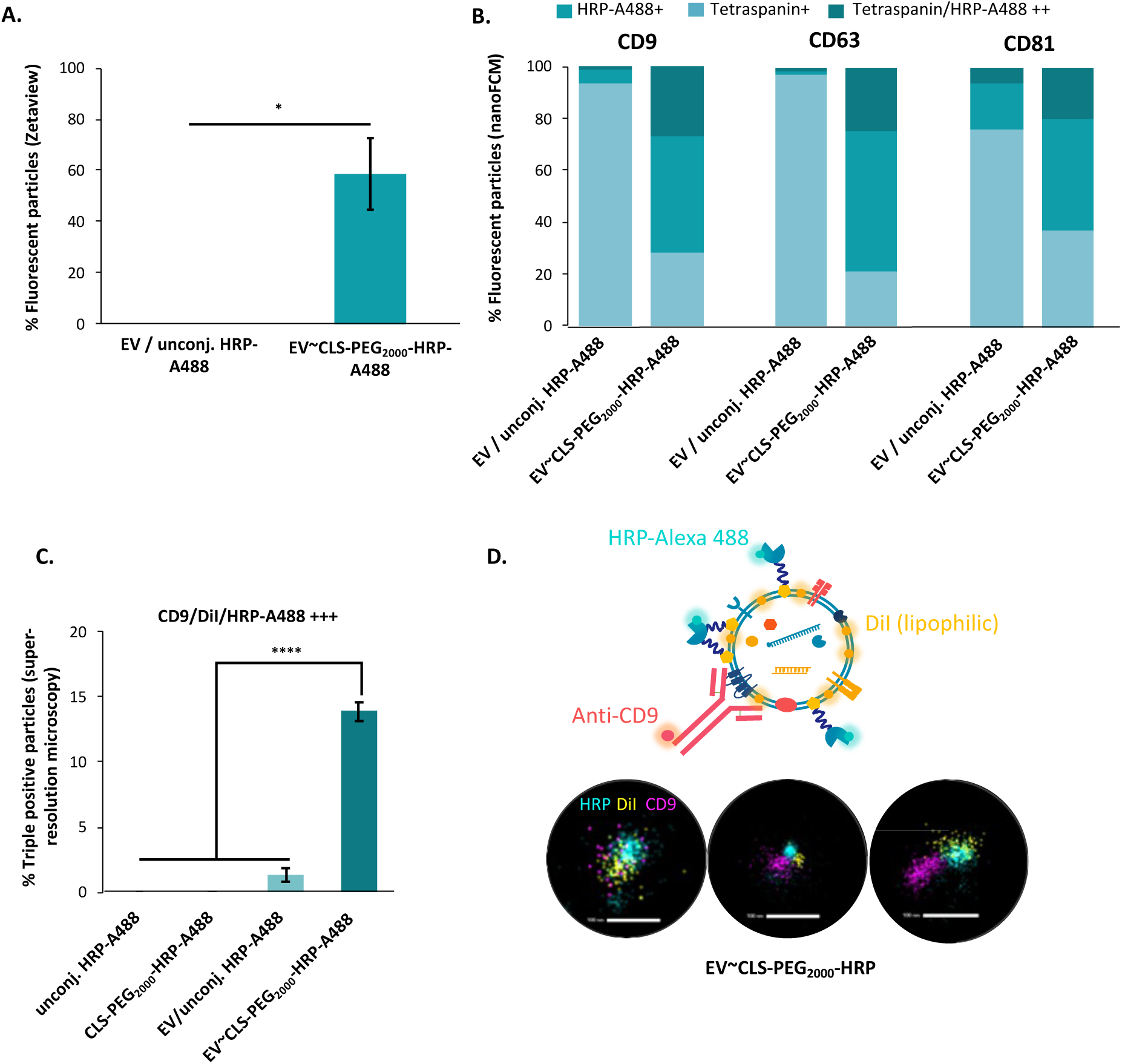
Heterogeneity in CLS-PEG_2000_-HRP-A488 loading. **(A)** Proportion of fluorescent detected particles using Zetaview (n=3) **(B)** NanoFCM analysis after incubation with anti-CD9, anti-CD63 or anti-CD81 antibodies coupled with AlexaFluor647 (n=1). Only fluorescent particles have been considered. **(C)** Proportion of triple positive clusters using STORM. Samples were incubated with DiI dye and anti-CD9 antibody, while HRP was labeled with Alexa-Fluor 488 (n=3). **(D)** Representative images of triple positive clusters observed using STORM. p-values were calculated using a two-tailed Welch test. *n.s*>0.05, *<0.05, **<0.01, ***<0.001, ****<0.0001.

Then, NanoFlowCytoMetry (nanoFCM) analysis were carried out using Alexa647-coupled antibodies anti-CD9, -CD63 and -CD81, three EV markers (Figure 3B, details are provided in Supplementary Figure 4). We did not purify samples by SEC to reach an EV concentration compatible with nanoFCM sensitivity. No A488+ particles were detected in control buffer containing CLS-PEG_2000_-HRP-A488 alone (Supplementary Figure 4), demonstrating that no fluorescent CLS-PEG_2000_-HRP-A488 aggregates were present. This control rules out the possibility that such aggregates of HRP could be responsible for positive artifacts in EV samples. Overall, 72%, 79% and 63% of fluorescent particles were A488+ in EV∼CLS-PEG_2000_-HRP-A488 after incubation with anti-CD9, -CD63 and -CD81 antibodies respectively. These values were 7%, 2% and 24% for EV / unconjugated HRP-A488. 27%, 24% and 20% of fluorescent particles were double positives for A488 and respectively CD9, CD63 and CD81 in EVs∼CLS-PEG_2000_-HRP-A488 sample in contrast to EV / unconjugated HRP-A488 showing respectively 1%, 1% and 6% of double positive EVs.

Finally, we used super-resolution microscopy (STORM) observations on EVs labeled with DiI lipophilic dye and anti-CD9 antibody (Figures 3C&D). This last technology enabled the observation of the colocalization of HRP with EVs, both qualitatively and quantitatively. In contrast to controls, 14±1% of EV∼CLS-PEG_2000_-HRP-A488 particles were triple positive. Details in number of cluster-types are provided in Supplementary Figure 5.

We also tested a second EV product to investigate if this relatively low proportion of triple-positive EV was due to the EV origin and to control the versatility of this approach. These EVs were produced from mMSC under 2D-culture condition and purified by ultracentrifugation (characterization is shown in Supplementary Figure 6). 42±2% of these EVs were triple positive after association with CLS-PEG_2000_-HRP-A488 (Supplementary Figure 7). This could be due to differences in membrane composition and tetraspanin expression. Overall, these results highlight the versatility of this approach to associate a protein to different EVs and demonstrate a high engineering efficiency approximatively equal to 60%.

#### 2.1.4. Lipid-PEG-cargo association does not modify EV size and surface characteristics

Then, we investigated the potential impact of LA conjugation on physico-chemical properties of EVs since proteins are large cargos that could disturb EV membranes. EV∼CLS-PEG_2000_-HRP were analyzed by NTA for size (Figure 4A) and Zeta Potential (ZP, Figure 4B). We did not observe significant changes neither in mode (110±4nm for EV∼CLS-PEG_2000_-HRP and 112±5nm for control) nor in negative ZP (−20±1mV for EV∼CLS-PEG_2000_-HRP and -19±1mV for control) after association. We compared EV / unconjugated HRP and EV∼CLS-PEG_2000_-HRP using MACSPlex assay to evaluate the impact of the association on EVs antigens accessibility. Interestingly, EV∼CLS-PEG_2000_-HRP and control exhibited similar profiles (Figure 4C). 33 EV surface epitopes assessed by the test were detected in both types of EVs highlighting the preservation of the accessibility of EV surface antigens using this association method. CryoTEM imaging revealed no difference between EV samples associated or not with CLS-PEG2000-HRP (Figure 4D).

**FIGURE 4.**
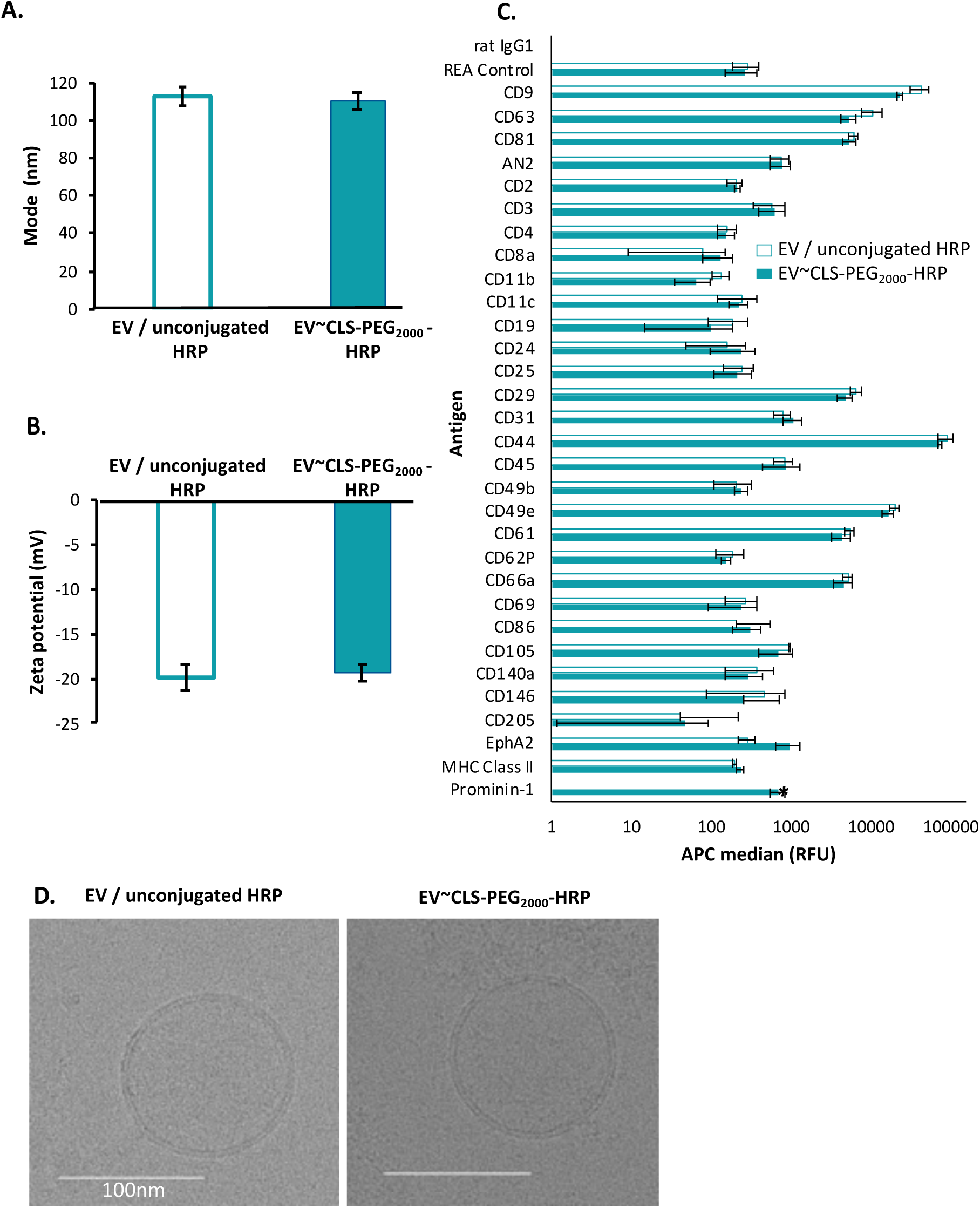
Characteristics of mMSC-EV associated or not with Lipid-PEG-HRP. **(A)** Size (mode, NTA, NS300, n=5) and **(B)** Zeta potential (NTA, Zetaview Twin, n=3) after incubation of EVs with unconjugated HRP or CLS-PEG_2000_-HRP and SEC separation. **(C)** MACSPlex Assay analysis as APC median after subtraction of buffer background signal. p-values were calculated using a two-tailed Welch test and only Prominin-1 was significant (*p-value<0.05). **(D)** CryoTEM images of EVs after incubation with unconjugated HRP or CLS-PEG_2000_-HRP. No SEC separation was processed on these samples (n=3).

#### 2.1.5. Lipid-PEG-cargo anchorage to EVs demonstrates a good stability in presence of serum

A common reported issue using LA conjugation to EVs is the redistribution of the cargo to plasma components as described in some studies with small cargos^29,38^. For comparison purpose, we carried out similar stability study on EV∼Lipid-PEG_2000_-HRP with DSPE and CLS lipid. For the DSPE lipid (Figure 5A), 107±34%, 94±13%, 106±15% and 65±3% of the associated protein remained with EVs after 4°C ON in DPBS, a freeze/thaw (F/T) in DPBS, 37°C during 2 hours in DPBS or DPBS supplemented with Fetal Bovine Serum (FBS, 1:50), respectively. For CLS lipid (Figure 5B), these values were 75±16%, 159±52%, 83±14% and 71±1%, respectively. Overall, these results showed a good stability of the anchorage with EVs despite some partial redistribution of Lipid-PEG-HRP to FBS components that seems to be impacted by the lipid nature of LA.

**FIGURE 5.**
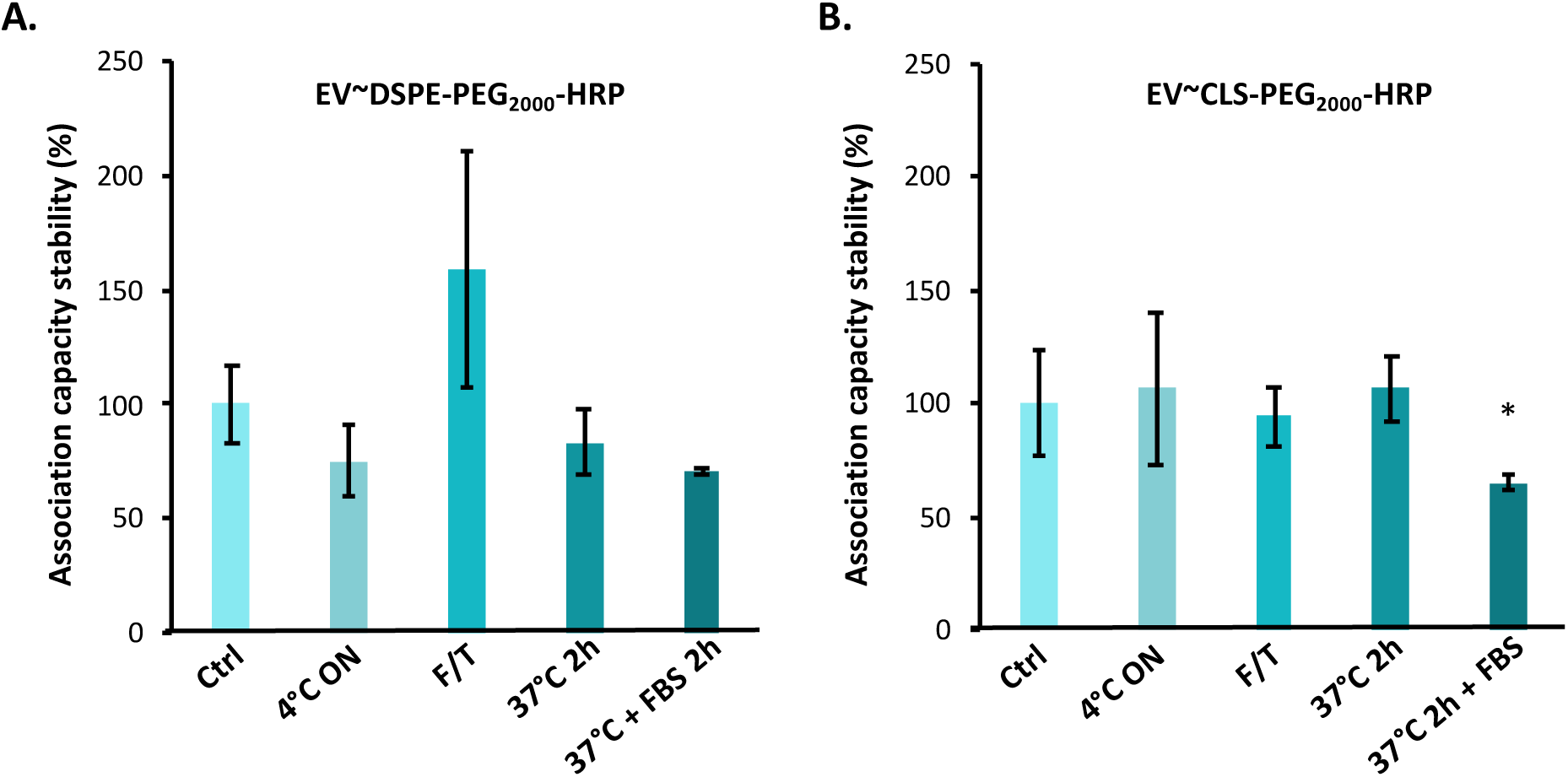
Stability of the EV / Lipid-PEG_2000_-HRP association. Association capacity of **(A)** DSPE-PEG_2000_-HRP and **(B)** CLS-PEG_2000_-HRP in percentage of control samples. Samples were diluted at 1:11 in DPBS (or DPBS supplemented with 1:50 FBS if indicated) and incubated before SEC separation. **4°C ON** samples were incubated at 4°C ON, **F/T** were frozen at −80°C before thawing, and **37°C 2H (+FBS)** were incubated at 37°C during 2h either with DPBS or DPBS + 1:50 FBS. n=3 except for CLS-PEG_2000_-HRP 4°C ON (n=2). p-values were calculated using a two-tailed Welch test and samples were compared with Ctrl. n.s>0.05, *<0.05.

### 2.2. Control of EV / LA-protein association process enables its efficient and functional delivery into carcinoma cell lines

#### 2.2.1. EV association enables concentration-, time-and cell-dependent intracellular delivery of active cargo protein into PANC-1, A549 and SKBR3 cells

After optimizing LA-protein cargo association to EVs to obtain high loading close to saturation, we aimed at demonstrating that this strategy was able to turn EVs into effective protein delivery systems by characterizing the intracellular delivery with absolute amount of vectorized cargo. EV∼Lipid-PEG-HRP samples were incubated with 1.10^5^ PANC-1 cells (adherent epithelioid carcinoma cell line derived from human pancreas) per well. Cells were subsequently washed before lysis and HRP enzymatic activity was measured in cell lysate. Importantly, we carried out all these experiments in presence of FBS in order to verify the stability of the anchorage suggested by the previous stability experiment. Number of EVs per cell was variable between experiments depending on the EV/HRP association capacity but remained in the range of 500-2000 EVs/cell to still in physiologically relevant conditions^39^.

First, EV∼CLS-PEG_2000_-HRP were tested for cell internalization at different HRP amounts (1ng, 5ng, 10ng and 15ng per well, Figure 6A). Native EVs without HRP did not trigger any peroxidase activity in PANC-1 cells lysate, validating the absence of detectable endogenous peroxidase activity. We observed a slight increase in HRP activity when cells were treated with free CLS-PEG_2000_-HRP control with up to 64±9pg recovered at 15ng incubated. By contrast, we observed a significant increase in HRP activity when cells were incubated with EV∼CLS-PEG_2000_-HRP: from 18±2pg at 1ng/well to 310±50pg at 15ng/well incubated (Figure 6A). Incubation time impact on HRP internalization was assessed after 2h, 4h and 24h for an amount of HRP fixed at 10ng/well (Figure 6B). Internalized HRP increased over time from 14±2pg after 2h to 27±4pg after 24h for controls and from 56±5pg after 2h to 194±27pg after 24h for EV∼CLS-PEG_2000_-HRP. Then, we investigated the mechanism of this internalization by inhibiting it at 4°C (Figure 6C). Only 4±1pg and 15±4pg of HRP were recovered at 4°C compared to 8±2pg and 97±12pg at 37°C for CLS-PEG_2000_-HRP control and EV∼CLS-PEG_2000_-HRP respectively, suggesting a temperature-dependent internalization process as expected from endocytosis. Finally, we tested two other recipient cells lines in addition to PANC-1 cells, namely A549 and SKBR3 cells (adherent epithelioid carcinoma cell line derived from human lung and breast, respectively), to evaluate the versatility in protein delivery of this approach for other cell lines (Figure 6D). 358±16pg, 230±21pg and 119±55pg of HRP were recovered in cells lysates of A549, PANC-1 and SKBR3 cells respectively, demonstrating the ability of EV∼CLS-PEG_2000_-HRP to be internalized in different human adherent epithelioid carcinoma cell line *in vitro*. We also validated the versatility of this approach by using the second source of mMSC-EVs (previously described) to associate and deliver HRP to PANC-1 cells (Supplementary Figure 8).

**FIGURE 6.**
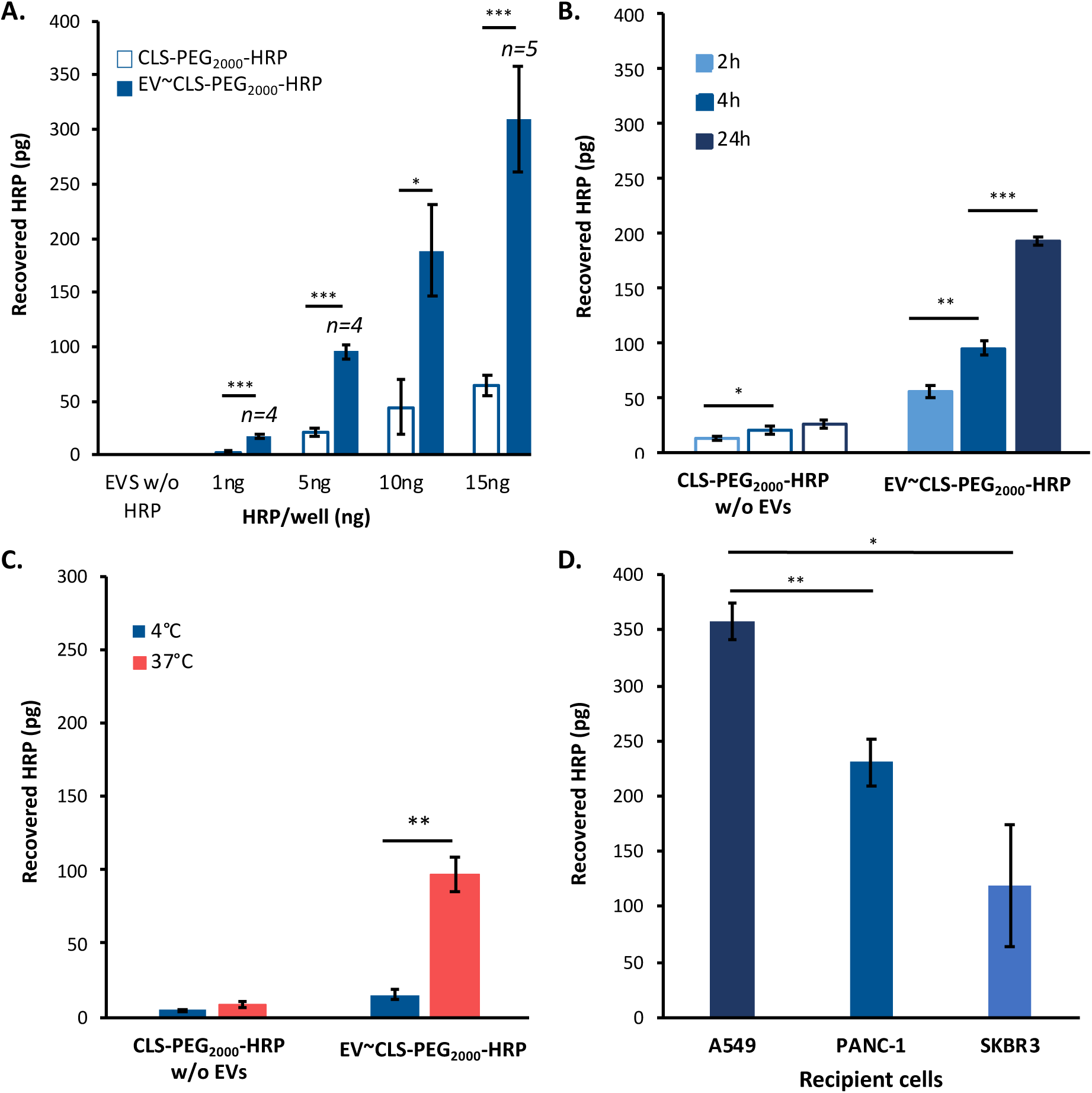
Quantification of absolute amount of internalized HRP depending on quantity, temperature and incubation time on cell internalization of EV∼CLS-PEG_2000_-HRP. Recovered CLS-PEG_2000_-HRP vectorized or not by EVs in PANC-1 lysate after incubation with 1.10^5^ PANC-1 following these conditions: **(A)** 4h, 37°C, **1ng, 5ng, 10ng or 15ng** of HRP/well. **(B) 2h, 4h or 24h** of incubation, 37°C, 5ng/well of HRP. **(C)** 2h, **4°C or 37°C**, 10ng/well of HRP. **(D)** 4h, 37°C, 10ng with 1.10^5^ **A549, PANC-1 or SKBR3 cells**. For each experiment, p-values were calculated using a two-tailed Welch test. *n.s*>0.05, *<0.05, **<0.01, ***<0.001. n=3 if unspecified.

In order to control and validate the intracellular location of anchored cargo after delivery, A549 cell internalization was observed using both epifluorescence microscopy (Figure 7A to 7E) and confocal microscopy (Figure 7F). Fluorescent spots were only visible in EV∼CLS-PEG_2000_-HRP-A488 (Figure 7E) and not in controls. Due to the limited Alexa-Fluor labeling of the HRP (resulting from a limited number of free lysines after LA-conjugation), we analyzed 3D-stack images using a random-forest algorithm to determine the number of fluorescent spots inside cells (Figure 7F). We considered this approach to be more objective than “naked-eyes” observation of the stack, especially due to the expected location of this lipidated HRP to the inner cell membrane once delivered. 52±6 spots per 1.10^6^ voxels were detected in EV∼CLS-PEG_2000_-HRP-A488 in contrast to controls that presented <1 spot per 1.10^6^ voxels. Taken together, these results confirmed quantitative cell lysis experiments.

**FIGURE 7.**
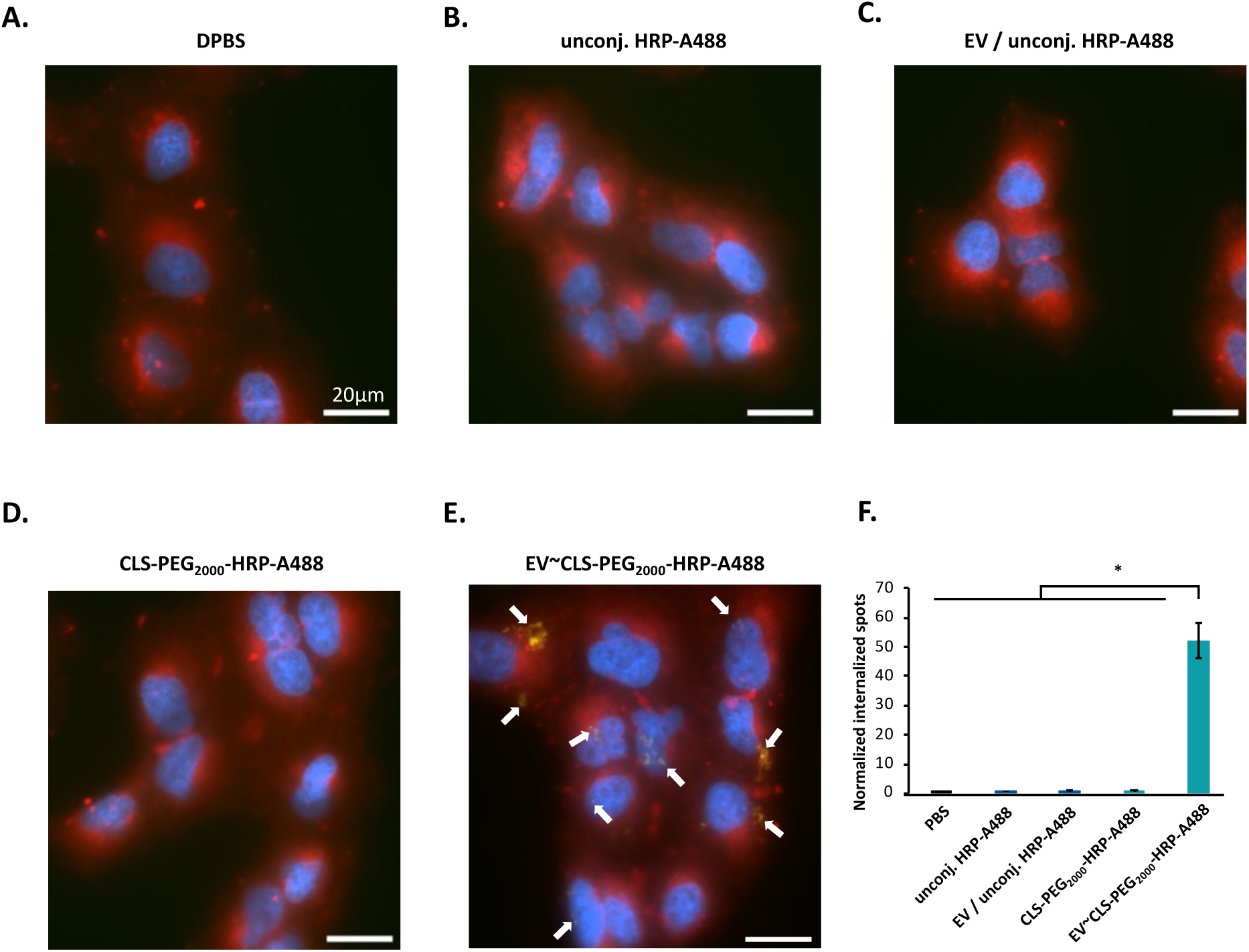
Epifluorescence and confocal microscopy observation and quantification of HRP-A488 cell internalization. A549 cells were incubated 4 hours with HRP-A488, conjugated with CLS-PEG_2000_ or not and associated with EVs or not. Panel images have been acquired using full-field epifluorescence microscopy. Blue, Red and green fluorescence are DAPI, Cell Mask lipophilic dye and Alexa-fluor 488 from labeled HRP, respectively. **(A)** A549 cells control incubated with DPBS. **(B)** and **(C)** A549 cells incubated with unconjugated HRP-A488 alone and mixed with EVs, respectively. **(D)** and **(E)** A549 cells incubated with CLS-PEG_2000_-HRP-A488 alone and associated with EVs, respectively. Identical HRP mass was deposited between EV conditions and their respective controls. Identical number of EVs was deposited between EV conditions. Bar scale is 20µm. White arrows highlight some HRP-A488 spots. **(F)** Quantitative analysis on confocal 3D image stacks obtained using spinning-disk confocal microscopy. Number of fluorescent spots in cells was normalized per total number of voxels corresponding to intracellular space. n=2 images except for EV / unconj. HRP-A488 (n=3). For each experiment, p-values were calculated using a two-tailed Welch test. *n.s*>0.05, *<0.05.

#### 2.2.2. Lipid-PEG linker MW impacts cell internalization of EVs∼Lipid-PEG-HRP by steric hindrance of native EV surface ligands

Finally, we evaluated impacts of lipid and PEG moieties of Lipid-PEG anchors on the cell internalization (Figure 8A). In contrast to controls, EV∼CLS-PEG_2000_-HRP, EV∼DSPE-PEG_1000_-HRP, EV∼DSPE-PEG_2000_-HRP and EV∼DSPE-PEG_5000_-HRP were internalized at 124±22pg, 140±8pg, 87±4pg and 12±5pg respectively. Regarding these results, it seemed that the PEG linker MW strongly influenced the cell internalization efficiency. We hypothesized that this could be due to steric hindrance and that a control of the density of DSPE-PEG_5000_-HRP per EVs could restore their internalization properties. To assess this assumption, we investigated the influence of a decreased density of cargos associated per EV on cell internalization (Figure 8B to 8D). Interestingly, only EV∼DSPE-PEG_5000_-HRP showed improvements in cell internalization by 1.4±0.1 and 2.1±0.3 fold-change thanks to a decreased density from 337 to 171 and 84 HRP/EV. Lack of changes in EV∼CLS-PEG_2000_-HRP and EV∼DSPE-PEG_1000_-HRP strongly suggests that steric hindrance of PEG_5000_ was responsible for this observation, underlining the interest to control these parameters to maintain EV / target cells interactions.

**FIGURE 8.**
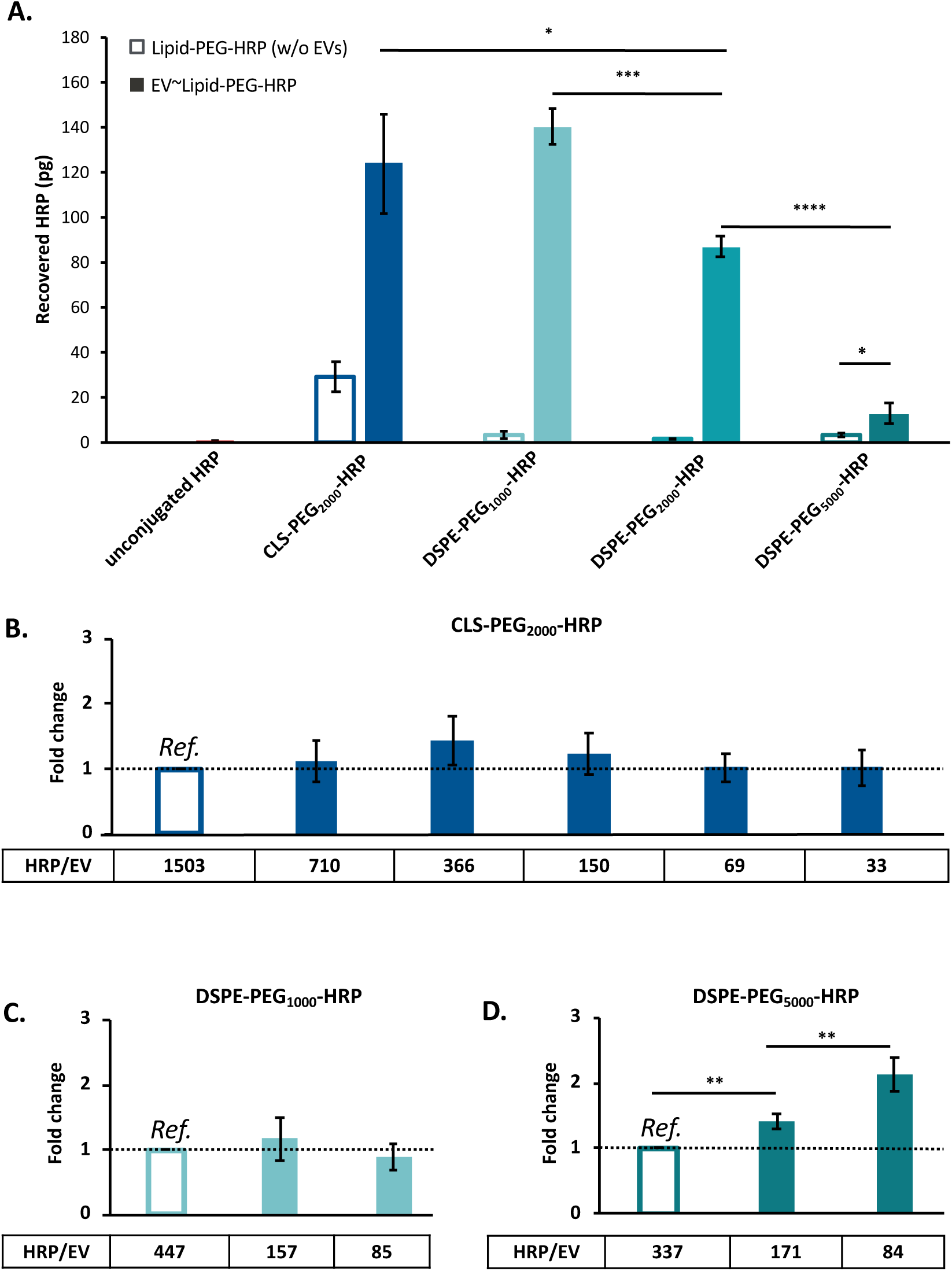
Impact of lipid anchor nature, PEG linker MW and HRP/EV ratio on cell internalization. **(A)** Recovered Lipid-PEG-HRP in PANC-1 lysate after incubation of 10µg of Lipid-PEG-HRP associated or not to EVs, for 4h at 37°C with 1.10^5^ PANC-1 cells (n=4). **(B), (C) and (D).** Fold change in recovered Lipid-PEG-HRP in PANC-1 lysate after incubation of 1500 EV∼Lipid-PEG-HRP per cell for 4h at 37°C with 1.10^5^ PANC-1 cells. Decreasing HRP/EV ratio have been tested with CLS-PEG_2000_-HRP, DSPE-PEG_1000_-HRP and DSPE-PEG_5000_-HRP for **(B)**, **(C)** and **(D)**, respectively. Proportion of Lipid-PEG-HRP recovered in lysate was normalized with the highest ratio of the experiment (indicated on figure as *Ref.*). Significances are compared to 1 (null hypothesis = no changes whatever the HRP/EV ratio) (n=3). For each experiment, p-values were calculated using a two-tailed Welch test. *n.s*>0.05, *<0.05, **<0.01, ***<0.001, ****<0.0001.

## 3. Discussion

### Lipid-anchorage is a versatile approach compatible with protein cargos while maintaining their activity

The main aim of this study was to study fundamental parameters controlling EV / LA-protein association to provide a versatile way to use EVs as effective delivery systems for intracellular vectorization of a functional protein. Indeed, no cellular internalization of an active protein associated with EVs using LA-based approach has been reported for now. We chose to use a click-chemistry reaction between maleimide and thiol to associate Lipid-PEG and cargo protein to take advantage of the high versatility of this approach, compatible with virtually all cargo proteins. While no unconjugated cargo was loaded by simple incubation, up to 898±247 cargos were associated per EV (Figure 2B). Assuming that HRP cargo and EV hydrodynamic radius are egal to 3nm^40^ and 50nm, respectively, one can estimate an EV surface coverage of ≈90% (using the *smoothed octagon packing onto a sphere* model^41,42^. This calculation neglects EVs larger than 100nm and the EV surface fraction already occupied by native membrane proteins but suggests that LA anchorage enables a very high association efficiency. Interestingly, this coverage density did not prevent EV/cell interaction using PEG < 5000g/mol, since decrease in density using medium PEG had no effect (Figure 8B and 8C).

By optimizing the amount of cargo protein during incubation, we reached an association efficiency of 53% (after correction of the measured value which was underestimated due to the loss of EVs during SEC) (Figure 2C), which is among best protein loading efficiencies with EV reported into the literature^27^. We used three different single-particle analysis methods to characterize association heterogeneity, namely fluorescent NTA, nano-flow cytometry and super-resolution microscopy. Overall, up to 70% of particles were cargo+ (Figures 3A&B), among which half were tetraspanins+ (Figure 3B) according to nanoFCM measurements. This supports observations already done by other groups^38,43^ and highlights the importance to characterize the loading/association efficiency along with other metrics (i.e., association capacity and efficiency). However, it seems important to underline than our labeling strategy was experimentally limited because majority of HRP lysines was already used for the LA-conjugation, which left few free lysines for the labeling with Alexa-Fluor. This has likely resulted in the underestimation of HRP positive EVs. Moreover, it has been demonstrated that antibody labeling of EVs could have a limited efficiency ^44^ and that all EVs do not express the three targeted tetraspanins (CD9, CD63 and CD81)^45,46^

Versatility of this approach has been controlled with another EV product. These EVs were also produced by mMSC but produced in 2D flask culture and isolated using ultracentrifugation (in contrast to the main commercial source of EVs used in this work, produced under turbulent flow from mMSC seeded onto microcarriers and isolated using SEC and tangential flow filtration). These EVs presented a different membrane composition in terms of surface antigens (Supplementary Figure 6). We observed differences in association efficiency (Supplementary Figure 7), capacity and cell delivery (Supplementary Figure 8A and 8B, respectively) which could be due to differences in the corona density and/or composition along with the membrane lipid content. Such impact of the EV source of LA-conjugation has been reported by comparing EVs from different cells sources^29^ but the impact of the production and purification process on LA-conjugation was not reported, to the best of our knowledge.

We showed that cholesterol (CLS) was 3.5-fold more efficient for association than DSPE with mMSC derived EVs (Figure 2B). This greater affinity of CLS could be due to the cholesterol condensing effect^47^ combined to its natural presence in high amount in EV membrane^48^. Supporting this assumption, a recent study^38^ demonstrated the ability of EV membrane to insert up to 20% exogenous cholesterol in addition to endogenous cholesterol, instead of replacing it. Interestingly, this outperformance by the cholesterol anchor has also been observed for siRNA cargos^33,37^ but not by some other teams working with smaller chemical cargos^29^. These results suggest that the presence of a large protein cargo did not reduce CLS anchoring efficiency.

### Control of Lipid Anchorage enables intracellular delivery of cargo protein and is affected by LA structure

After successful association study and optimization, we investigated intracellular delivery of cargo protein. Regarding these results, CLS-PEG_2000_ and a ≈1000 cargo/EV ratio were chosen as standard conditions. This high number of cargo associated per EVs obtained in this study enabled the use of low amount of EVs to treat cells, with 200 to 2000 EVs per cell, a dose compatible with cargo-specific cell response as recently established^39^. Even in presence of serum, association and vectorization of protein cargo with EVs enabled up to 50 folds more intracellular delivery in three different carcinoma-derived cell lines (PANC-1, A549 and SKBR3 cells). These results suggest that proteins could be more stable cargos for lipid anchorage than small chemicals^29,38^. Some differences in protein delivery were observed among cell types, confirming the already described phenomenon that EV internalization can differ depending on cell type^17^. Interestingly, this internalization was concentration-, time- and temperature-dependent, allowing for a fine tuning of the amount that can be delivered. Unfortunately, it is difficult to compare these masses of internalized cargo with literature since such absolute measurements are barely ever reported. Finally, we highlighted the impact of three PEG linker MW on cell internalization. Indeed, when the cargo protein was associated to EVs with a high MW PEG linker (5000g/mol), the cell internalization efficiency was significantly decreased. This effect was mitigated by decreasing the number and therefore density of DSPE-PEG_5000_-HRP per EV. Such impact of the LA linker on EV / cargo association and cellular uptake was never reported and should probably be considered in future studies.

## 4. Conclusion

While EVs are promising biomolecule vectors, their membrane is as impermeable to proteins as the cellular one. Thus, reaching acceptable protein loading capacity and intracellular delivery remains an important challenge. Altogether, we demonstrated that exogenous association of active proteins to EVs using LA is possible and can lead to efficient intracellular delivery systems, even in the presence of serum. By thoroughly characterizing this association and numerous governing factors (Lipid nature, PEG linker, temperature, EV origin, etc.), it was possible to control the number of HRP associated to EV surface, allowing for efficient and tunable intracellular delivery. These results prove the particular interest to thoroughly characterize EV-association processes with the aim to preserve their natural properties for cell interactions. To improve this process, other molecules such as lipids and polymers could be investigated. In particular, the effect of the PEG linker has been by far less studied than lipid impact whatever the type of cargo, despite being crucial as showed in this study. Thus, this study highlights the value of controlled and rational engineering of EVs to make them efficient biomolecule delivery systems, opening up promising prospects for various protein replacement or inhibition therapies.

## 5. Materials and methods

### 5.1. Chemicals

Chemicals were purchased from (i) Biopharma PEG (USA): DSPE-PEG_1000_-SH, DSPE-PEG_2000_-SH, DSPE-PEG_5000_-SH, CLS-PEG_2000_-SH (structures are shown in Supplementary Figure 1A); (ii) Sigma-Aldrich (France): sepharose CL-2B cross-linked, sulfuric acid 95-97%; formic acid 98-100%; acetonitrile 99%, trehalose, Tris-HCl; (iii) Thermofisher (France): EZ-Link™ Maleimide Activated Horseradish Peroxidase, ELISA 1-Step™ Ultra TMB, M-PER™ Mammalian Protein Extraction Reagent, Pierce™ Protease Inhibitor Mini Tablets, CellMask™ Plasma Membrane Stains, DAPI, ProLong™ Gold Antifade Mountant; (iv) Gibco LifeTechologies (USA): Dulbecco’s Modified Eagle Medium (DMEM), Foetal Bovine Serum (FBS), penicillin/streptomycin, L-glutamine, trypsin-EDTA, opti-MEM, Dulbecco’s Phosphate-Buffered Saline (DPBS).

### 5.2. Murine Mesenchymal stem cell derived Extracellular Vesicles (mMSC-EVs)

mMSC-EV were obtained from Everzom (Paris, France, products #230801-G5 and #230608-G12). Briefly, EVs from Murine C3H/10T1/2, Clone 8 (ATCC® CCL-226™) cells were produced and isolated using Everzom’s optimized 3D biomanufacturing and isolation processes, followed by an additional purification step with Size Exclusion Chromatography (SEC; Captocore 700). The column volume was adapted to the protein concentration measured after Tangential Flow Filtration (TFF). A 1:10 addition of 250mM trehalose (for a final concentration of 25mM) was done before freezing the sample in cryotubes. For some additional experiments, a second EV product was used. These mMCS-EV#2 were produced following a well-established protocol in our laboratory^49^. The final EV pellet was resuspended in DPBS buffer containing 25mM of trehalose and stored at −80°C.

### 5.3. Synthesis of HRP conjugates

#### 5.3.1. Synthesis of Lipid-PEG-HRP

Stock solutions of Lipid-PEG-SH (structures and conjugation process are described in Supplementary Figure 1A&B) were obtained by dissolving 20g/l of powder in ultrapure water. EZ-Link™ Maleimide Activated Horseradish Peroxidase (mal-HRP) was rehydrated in H_2_O for a final concentration of 2g/l before addition of 20 molar equivalents of Lipid-PEG-SH (Lipid-Anchor abbreviated as LA). Depending on the LA MW and the final volume needed, 45µl to 122µl of lipid-PEG-SH were added to 500µl to 1000µl of mal-HRP. Water was added according to the volume of added LA to obtain identical final concentration is all mixes, equal to 7.8×10^−4^M of Lipid-PEG-SH (1.4 to 4.5g/l according to the MW of the LA) with 3.9×10^−5^M (1.6g/l) of mal-HRP. The mix was then incubated at 37°C overnight under gentle agitation. A dialysis step (Slide-A-Lyzer^®^ Dialysis Cassette 20000 MWCO, 0.5ml, Thermo Fisher Scientific®) was performed against DPBS under gentle agitation to eliminate unconjugated Lipid-PEG-SH. Dialysis buffer was frequently changed (2h at RT, 2h at 30°C and finally 20h at 4°C). MW of DSPE-PEG_1000_-HRP, DSPE-PEG_2000_-HRP, DSPE-PEG_5000_-HRP and CLS-PEG_2000_-HRP were observed using HPLC-MS (Supplementary Figure 2C) and mass gains of respectively 1392g/mol, 2362g/mol, 4815g/mol and 2154g/mol as compared to unconjugated HRP were measured. Enzymatic assay revealed that Lipid-PEG-HRP retained 80-95% of its biological activity regardless of the type of Lipid-PEG after conjugation and dialysis (Supplementary Figure 2D).

#### 5.3.2. Synthesis of fluorescent Lipid-PEG-HRP-A488

Alexa-Fluor 488 was conjugated with Lipid-PEG-HRP to obtain Lipid-PEG-HRP-A488 for single-particles analysis and cell internalization microscopic observations using Alexa Fluor® 488 Microscale Protein Labeling Kit (Thermo Fisher Scientific®) following manufacturer’s protocol. Fluorescent conjugates were purified using Zeba™ Dye and Biotin Removal Spin Columns (Thermo Fisher Scientific®) and 50mM of Tris-HCl were added to quench the reactivity of residual dye and avoid further artifactual conjugation of the dye with EV proteins.

### 5.4. Lipid-PEG-HRP and EVs association

5.10^10^ EV/ml (assessed by NanoTracking Analysis, NTA) were mixed with different final concentrations of Lipid-PEG-HRP (from 0.5mg/ml to 75mg/ml). Depending on the requirements of the experiment, volumes of preparation were comprised between 50µl and 1ml. Samples were incubated overnight at 25°C or 37°C depending on the experiment, under gentle agitation.

### 5.5. Purification of EV∼Lipid-PEG-HRP

SEC separation was used for all samples except in indicated microscopy experiments (Figure 8). Columns were packed following Tulkens *et al.*^50^ protocol. 50µl of EV/HRP mixture at 5.10^10^ EV/ml were diluted 11-fold in DPBS and 500µl of sample were deposited on the top of the column. 500µl elution fractions were collected and analyzed (NTA, enzymatic activity). Depending on the experiment and the purity level needed, only the 4^th^ fraction or the combination of 3^rd^, 4^th^ and 5^th^ fractions was collected. Samples used for microscopy observations in Figure 7 were purified using tangential ultrafiltration with Amicon® 100kDa 0.5ml (Merck Millipore®) in order to avoid the dilution resulting from SEC. 500µl of samples previously diluted by 10 in DPBS were deposited in Amicon® filter before centrifugation at 2000g during 15min at 4°C (Centrifuge 5804 R, Eppendorf®). Then, retentates were diluted by 10 in DPBS once again to obtain 500µl before a second identical centrifugation. Final EV-containing retentates were analyzed using NTA and TMB enzymatic assay.

### 5.6. Characterization

#### 5.6.1. NanoTracking Analysis (NTA)

EVs were analyzed with a NanoSight NS300 (Malvern Panalytical Instruments, UK). Samples were diluted with particle-free DPBS in order to obtain a concentration range between 1.10^8^ and 5.10^8^ particles/ml. Measurements were performed with a 405nm and a 488nm laser. Data were collected using the NanoSight NTA 3.4 software following a tailored script: temperature was set at 25°C, syringe pump at 40 arbitrary units and 5 videos of 60s were recorded with a camera level set to 15 and analyzed with a detection threshold set to 5. All groups of compared data were measured and analyzed using identical conditions. Zetaview® (Particle Metrix, USA) was used for Zeta potential (ZP) and particle fluorescence measurements. For ZP measurements, 2-cycles measurements on 2 positions (SL1 and SL2) using a 488nm laser were used with a sensitivity set at 80 and a shutter at 100. Samples were diluted to obtain a final EV concentration equal to 5.10^7^ EV/ml in 1:10 DPBS (1.5mM). Sensed conductivity was equal to 145µS/cm. For particle fluorescence measurement, two successive measurements were done in fluorescent and scatter mode. Fluorescence measurements were performed with a sensitivity of 90 and a shutter of 50 using a 488nm laser and a 500nm filter, while scatter measurements were carried out with a sensitivity of 80 and a shutter of 100 without filter. Concentration correction factor was calculated following manufacturer’s guidelines by using reference fluorescent beads YG-488 and was equal to 1.1. Data were analyzed using Software ZetaView (8.05.16 SP3 version).

#### 5.6.2. HPLC-MS experiments

ElectroSpray Ionization Mass Spectrometry (ESI/MS) and MS/MS analyses were performed on a Synapt G2-S instrument equipped with Ultra Performance Liquid Chromatography (Waters Corporation, UK). For the separation, a gradient with H_2_O containing 0.1% of formic acid (FA) as eluent A and acetonitrile containing 0.1% of FA as eluent B was applied as follow: from 95% A to 70% in 8min and from 70% until 50% between 8 and 25min before going back to 95% A in a total run of 35min, at a flow rate of 0.45ml/min on a polyphenyl column (BioResolve, 150×2.1mm, 2.7µm particle size), designed for intact proteins and antibodies analysis. UV detection was done at 220nm and 280nm. Mass spectrum acquisition was performed with a mass range of m/z 800-5000, a scan time of 0.5s in a positive ion mode with capillary voltage 3kV, 40°C and 350°C for the source and desolvation temperatures, respectively. MassLynx software was used for data acquisition and processing, with the MaxEnt1 module for the deconvolution process.

#### 5.6.3. Enzymatic assay and determination of EV-associated HRP amount

The amount of EV-associated HRP was measured by enzymatic activity using tetramethylbenzidine (TMB) after SEC separation. For each SEC-purified sample, an individual standard curve was generated using the pre-SEC sample (of known HRP concentration) to account for any possible variation due to the LA-HRP lot. 50µl of sample were added to 50µl of TMB and incubated 30min at RT in 96-well plate (Nunc MaxiSorp™, Invitrogen™). Then, 50µl of sulfuric acid (2M) were added to stop reaction. Absorbance at 450nm was then measured to quantify the HRP activity using a MultiScan Go (Thermo Fisher Scientific™). The association capacity was calculated by dividing the HRP concentration (expressed as the number of proteins/mL) by the EV concentration (number of particles/mL) and is expressed as the number of HRP/EV. Association efficiency was defined as the proportion of HRP remaining with EVs after SEC as compared to the initial amount during incubation with EVs and before SEC.

#### 5.6.4. MACSPlex assay

EV surface epitopes determination was performed using the MACSPlex Exosome Kit mouse (Miltenyi Biotec®). The test was done according to the manufacturer protocol. Briefly, 1.10^9^ EVs were diluted in 120µl MACSPlex Buffer. Detection cocktail [MACSPlex Exosome Capture Beads and Detection Reagent CD9, CD63 and CD81] was then added to the EV suspension and incubated for 1h at RT under gentle agitation (450rpm) protected from light. Capture beads contained 39 populations of dyed beads coated with monoclonal antibodies able to recognize 37 potential EV surface antigens along with 2 internal isotype negative controls. MACSPlex Exosome Detection reagent contained APC-labeled detection antibodies against the three tetraspanins CD9, CD63, and CD81. After washing steps, samples were analyzed using a BD Accuri™ C6 (BD Biosciences, USA, (software version 1.0.34.1 Build 20180111.34.1). For each sample, 7000–12,000 single-bead events were measured. Bead populations were gated based on their fluorescence intensity according to the manufacturer’s guidelines. The obtained median fluorescence intensity (MFI) of 37 exosomal surface epitopes and two isotype controls were subtracted from the corresponding MFI of the blank. This blank was the EVs buffer diluted in the same way as EV samples, mixed with capture beads and detection reagent without EVs.

#### 5.6.5. NanoFlowCytometry (nanoFCM)

NanoFCM™ was used in 40-200nm range mode. Alexa-Fluor 647 anti-mouse IgG1 (isotype control), APC anti-mouse CD63 Alexa-Fluor 647, APC anti-mouse CD9 Alexa-Fluor 647 and APC anti-mouse/rat CD81 Alexa-Fluor 647 (Biolegend) were diluted at 1:50000 before incubation with sample diluted at 5.10^8^ EV/ml at 4°C overnight (ON).

#### 5.6.6. Cryo-Transmission Electron Microscopy (cryo-TEM) imaging

Samples were imaged using cryo-TEM. Carbon film coated EM grids (Ted Pella Inc.) were glow-discharged at 30mA, for 30s (Pelco easiGlow, Ted Pella Inc.). 4μl of each sample solution was applied to the carbon-coated side of the TEM grid, blotted for 10s, before plunge-freezing in liquid ethane using the Vitrobot Mark MkIII (FEI). Vitrified samples were then imaged in a Glacios cryo-transmission electron microscope (ThermoFisher) equipped with a K3 direct electron detector, SerialEM software (D. Mastronarde, Boulder Lab) was used in acquisition.

#### 5.6.7. Super-resolution microscopy imaging

Single molecule localization microscopy was performed using an ONI Nanoimager (ONI Bio) following manufacturer guidelines. Native EV samples were incubated with CD9 Alexa-Fluor 488, CD63 Alexa-Fluor 568, and CD81 Alexa-Fluor 647 monoclonal antibodies from the EV Profiler kit (ONI Bio). EV∼LA-HRP samples were incubated with CD9 Alexa-Fluor 647 (Clone MZ3, BioLegend) and Vybrant DiI (Invitrogen, Thermo Fisher). Unbound antibody and dye were removed after overnight incubation using SEC (70nm qEV, IZON). 10µl droplet was added between a 1mm TOMO adhesion glass slide (Avantor) and a #1 glass cover slip (Corning) for immediate imaging. Briefly, the instrument was calibrated for direct stochastic optical reconstruction microscopy with bead calibration (ONI Bio) slide using NimOS version 1.19.7. 3-color super resolution microscopy was then performed with 500 frames taken per channel. Laser powers were adjusted according to company standard operating procedures. Raw image files were uploaded to the cloud software CODI via CODI Desktop Uploader version 0.30.0. Image analysis was performed on CODI. Briefly, drift correction, filtering, clustering, and counting algorithms were performed using the predefined “EV Profiling” workflow.

### 5.7. Cell internalization experiments

#### 5.7.1. Quantitative cell lysis experiments

1.10^5^ PANC-1 (ATCC, CRL-1469 ™), SKBR3 (ATCC, HTB-30™) and A549 (ATCC, CCL-185™) cells in 1ml of DMEM supplemented with 10% FBS, 2mM L-glutamine, 100U/ml penicillin and 100μg/ml streptomycin were plated in 24-well plate and incubated at 37°C and 5% CO_2_ prior to cell internalization experiments. After 24h, medium was discarded and replaced by samples to be tested diluted in fresh identical medium for a total volume of 400µl. Cells and samples were then incubated at 37°C with different amounts of HRP (from 1ng to 15ng of HRP/well), for different time periods (2h, 4h or 24h) depending on the experiment. Sample-containing medium was afterwards removed and cells were washed with cold DPBS 3 times before addition of 200µl of M-PER™ supplemented with protease inhibitor and incubated 5min at RT under gentle shaking. To ensure optimal lysis, cells were then pipetted in the well and collected in microtubes before 10s vortex. Lysates were finally centrifuged during 15min at 4°C and 14000g to keep only the supernatant. HRP activity was measured with TMB following protocol described in 5.6.3. except that all dilutions and standard curves were diluted in lysis buffer instead of DPBS to avoid any bias on the peroxidase activity from surfactants present in lysate. Blank was done with untreated cells control lysate.

#### 5.7.2. Microscopy observations and quantifications

A549 cells were seeded at 17500 cells per well in 8-well Nunc™ Lab-Tek™ II Chamber Slide™ plates (ThermoFischer) and incubated during 24h at 37 °C in a humidified 5% v/v CO_2_ atmosphere. Then, 150µl of samples diluted with 150µl of DMEM supplemented with 10% FBS, 2mM L-glutamine, 100U/ml penicillin and 100μg/ml streptomycin were added and incubated during 5h with cells. Cells were washed with DPBS, labeled with CellMask™ (1:1000 dilution in fresh medium) and incubated for 30min. Finally, cells were fixed in DPBS with 4% formaldehyde (36% w/w) during 10min at room temperature and washed with DPBS. DAPI (1µg/ml in DPBS) was added and left for 5min before washing. Slide was mounted with Prolong™ mounting medium for microscopy observations. For the panel (Figure 7A to 7E), images were acquired using a full-field epifluorescence microscope Nikon Ti2 (Nikon Instrument Inc®), with a CMOS back-illuminated Prime95B (1200*1200 pixels, 11µm pixel, Teledyne Photometrics®) camera, Lumencor® system as a source of illumination and 40X objective (oil, 1.3 numerical aperture). Images were treated with Fiji (ImageJ) software with identical adjustment of brightness and contrast. The set parameters minimum/maximum displayed values were equal to 3000/62000 for HRP-488, −140/17500 for CellMask™ and 6800/34500 for DAPI staining. For quantitative analysis, images were acquired on the same samples using a spinning-disk confocal microscope with a Yokogawa CSU-W1 spinning head (Yokogawa Electric Corporation®) on a sCMOS back-illuminated Prime95B (1200*1200, 11µm pixel) camera and Nikon 60X objective (Plan Apo, 1.4 numerical aperture). Images were taken with a Z-step of 210nm using an Oxxius® lasers (excitation at 488nm and 561nm). Image processing for quantitative analysis was performed using Fiji. Cells were segmented into a binary mask using the random-forest pixel-classifier algorithm (Labkit ImageJ plugin) to attribute a class (“inside cell” or “background”) to every pixel. Spot detection was done by applying a Laplacian of Gaussian filter before local minima detection (using FeatureJ Log filter and MorphoLibJ local minima detector) to extract a 3D point per spot. List of coordinates generated based on CellMask™ were used to differentiate cells from background in order to count HRP-A488 fluorescent spots inside cells. This number of internalized fluorescent spots was normalized per the total number of intracellular voxels in order to normalize the difference in cell number per image.

### 5.8. Statistical analysis

RStudio version 1.4.1717 (2009-2021 RStudio, PBC®) was used for *post hoc* statistical analysis. Unilateral or two-tailed Welch tests were used depending on the null hypothesis as indicated in figure legends.

## Acknowledgements

The authors gratefully acknowledge the French National Research Agency (ANR-20-CE09-0011-01), the imaging facility MRI, member of the France-BioImaging national infrastructure supported by the French National Research Agency (ANR-10-INBS-04, «Investments for the future»), IVETh Core Facility, and specifically Kelly Aubertin, Sarah Razafindrakoto and Christopher Ribes for fruitful discussions and for performing the NanoFCM analyses. IVETh is supported by the IdEx Université Paris Cité, ANR-18-IDEX-0001, by the Region Ile de France under the convention SESAME 2019 – IVETh (EX047011) and via the DIM BioConvS, by the Région Ile de France and Banque pour l’Investissement (BPI) under the convention Accompagnement et transformation des filières projet de recherche et développement N° DOS0154423/00 & DOS0154424/00, DOS0154426/00 & DOS0154427/00, and Agence Nationale de la Recherche through the program France 2030 “Integrateur biotherapie-bioproduction” (ANR-22-AIBB-0002). Funding for work by S. Emerson was provided by the California Institute of Regenerative Medicine Bridges program (EDUC2-12620).

## AUTHOR CONTRIBUTIONS

**Antonin Marquant**: Conceptualizaxon (lead); Invesxgaxon (lead); Methodology (lead); Data Curaxon; Formal analysis (lead); Validaxon; Visualizaxon; Wrixng—original dra{ (lead); Wrixng—review & edixng. **Jade Berthelot**: Conceptualizaxon; Invesxgaxon; Methodology; Validaxon, Vizualizaxon. **Claudia Bich**: Conceptualization; Invesxgaxon; Methodology; Validaxon, Vizualizaxon. **Zeineb Ibn Elfekih**: Invesxgaxon; Methodology; Validaxon; Vizualizaxon. **Laurianne Simon**: Conceptualization (supporting); Investigation. **Baptiste Robin**: Conceptualization (supporting); Investigation. **Joël Chopineau**: Conceptualization (supporting); Resources. **David Tianpei Wang**: Invesxgaxon; Methodology; Formal analysis; Data curaxon; Validaxon; Vizualizaxon. **Samuel J Emerson**: Invesxgaxon; Data Curaxon; Formal analysis. **Aijun Wang**: Conceptualization; Resources. **Clément Benedetti**: Invesxgaxon; Methodology; Formal analysis; Data curaxon; Validaxon; Vizualizaxon. **Simon Langlois**: Invesxgaxon; Methodology; Formal analysis; Data curaxon; Validaxon; Vizualizaxon. **Laurence Guglielmi**: Conceptualization (supporting); Methodology; Resources. **Pierre Martineau**: Conceptualization (supporting); Methodology; Resources. **Anne Aubert-Pouëssel**: Conceptualizaxon (lead); Supervision (co-lead); Visualizaxon; Wrixng—original dra{; Wrixng—review & edixng (co-lead). **Marie Morille**: Conceptualizaxon (lead); Supervision (lead); Validaxon; Vizualizaxon; Funding acquisixon; Project administraxon; Resources; Wrixng—original dra{ preparaxon; Wrixng—review & edixng (lead). **All authors** provided crixcal feedback and helped shape the research, analysis and manuscript.

## Supplementary Information

**Supplementary Figure 1.**
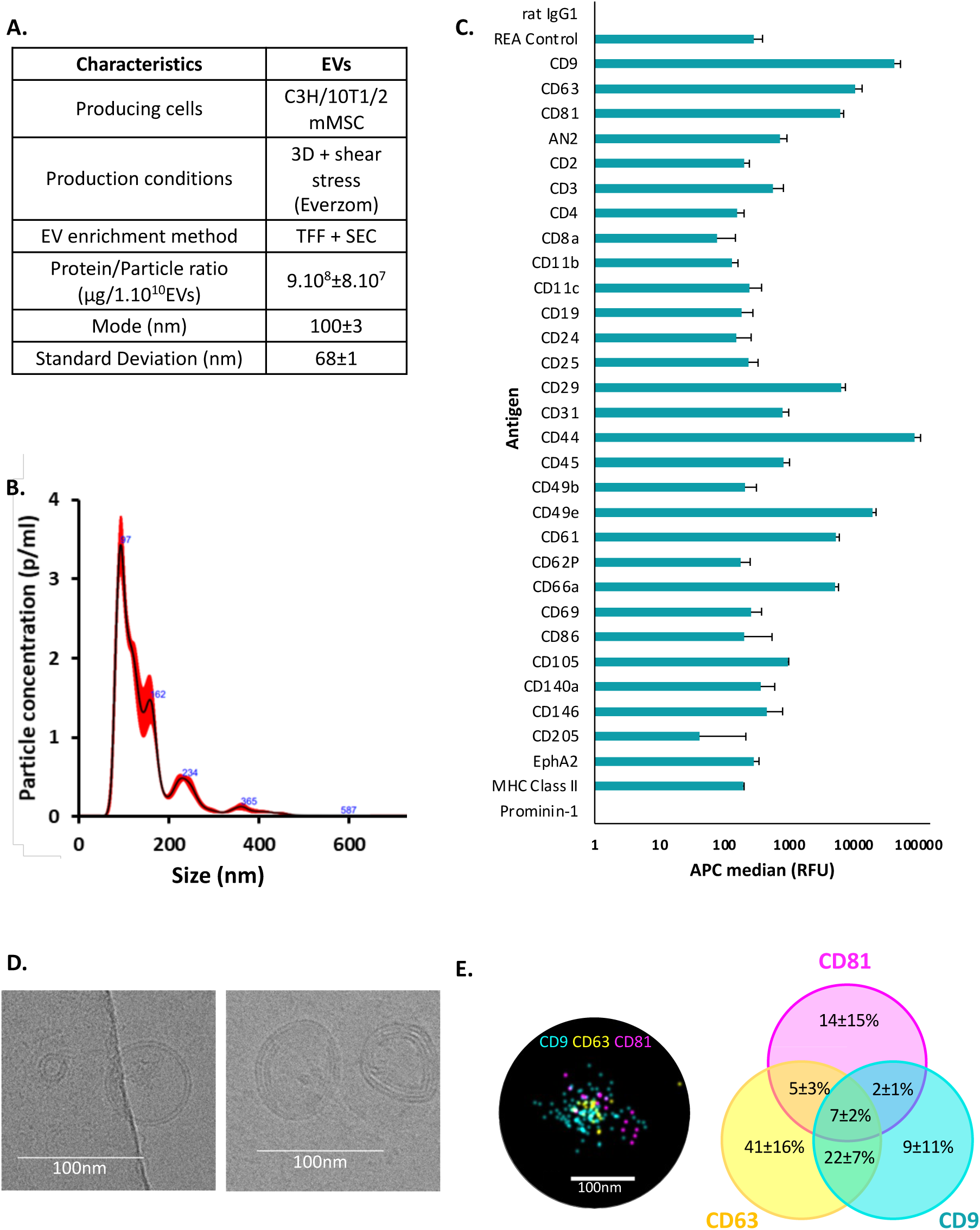
Characteristics of mMSC-EVs. **(A)** Production and enrichment characteristics. Protein full content was measured using µBCA assay and BSA as reference (n=3). EV concentration, mode and SD were measured using NTA (NS300). **(B)** Size distribution profiles measured by NS300. **(C)** MACSPlex Assay signal as APC median after subtraction of buffer background signal (n=3). **(D)** CryoTEM images. **(E)** Proportion of single, double and triple positive clusters analyzed using STORM super-resolution microscopy. Samples were incubated with anti-CD9, -CD63 and -CD81 antibodies. Representative images of triple positive clusters are shown. Bar scale is 100nm (n=3).

**Supplementary Figure 2.**
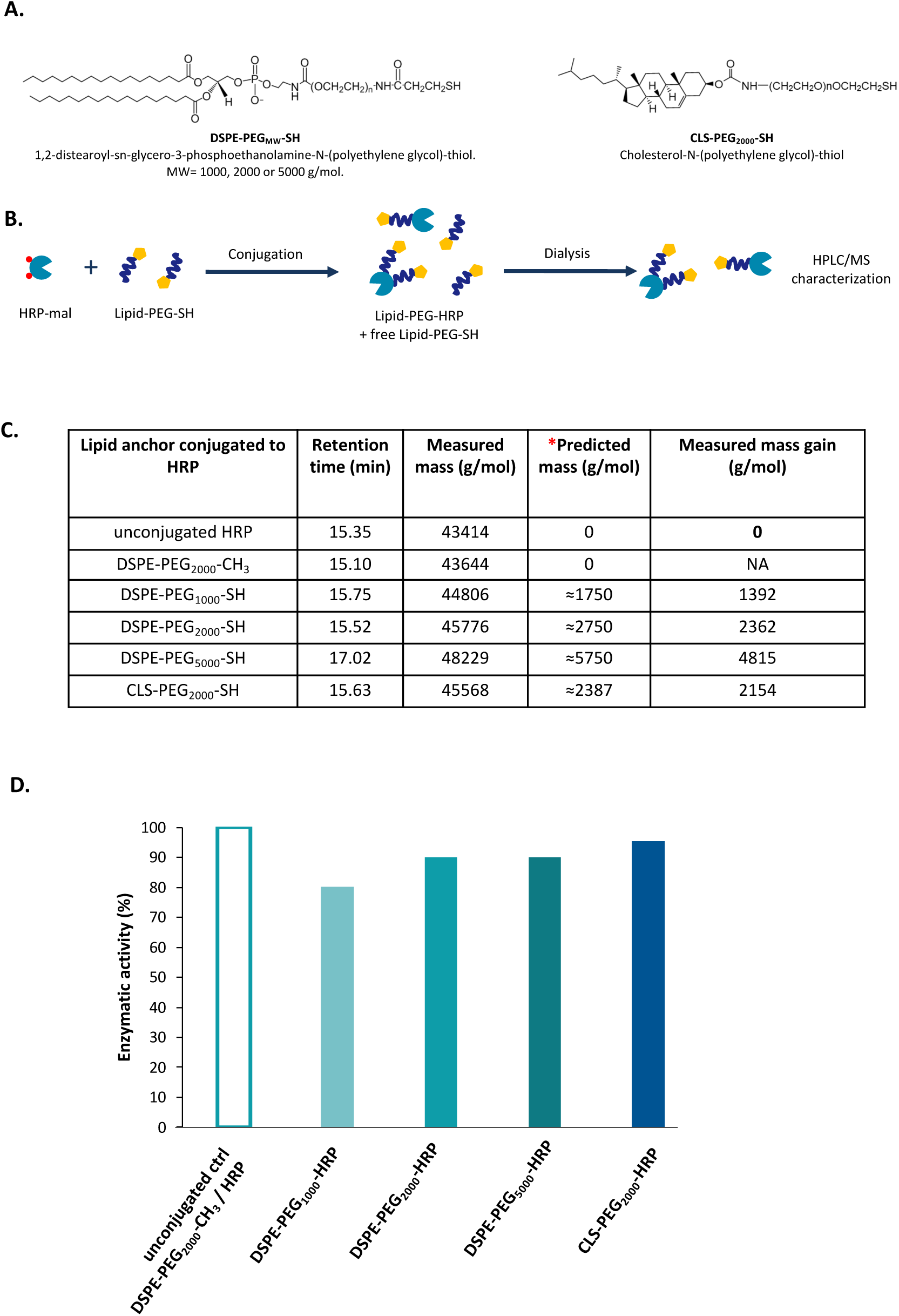
Characterization of Lipid-PEG-HRP. **(A)** Chemical structure of DSPE-PEG_MW_-SH (MW=1000, 2000 or 5000g/mol) and CLS-PEG_2000_-SH. **(B)** Scheme of Lipid-PEG-SH / mal-HRP conjugation process (not at scale). **(C)** HPLC-MS analysis of unconjugated HRP and Lipid-PEG-HRP. *Predicted mass are based on the mean MW of the PEG linker and the corresponding lipid anchor. **(D)** Enzymatic activity of Lipid-PEG-HRP after dialysis. Reference is unconjugated ctrl DSPE-PEG_2000_-CH_3_ / HRP to take into account of biased increase in HRP activity before dialysis due to the presence of lipid excess in samples.

**Supplementary Figure 3.**
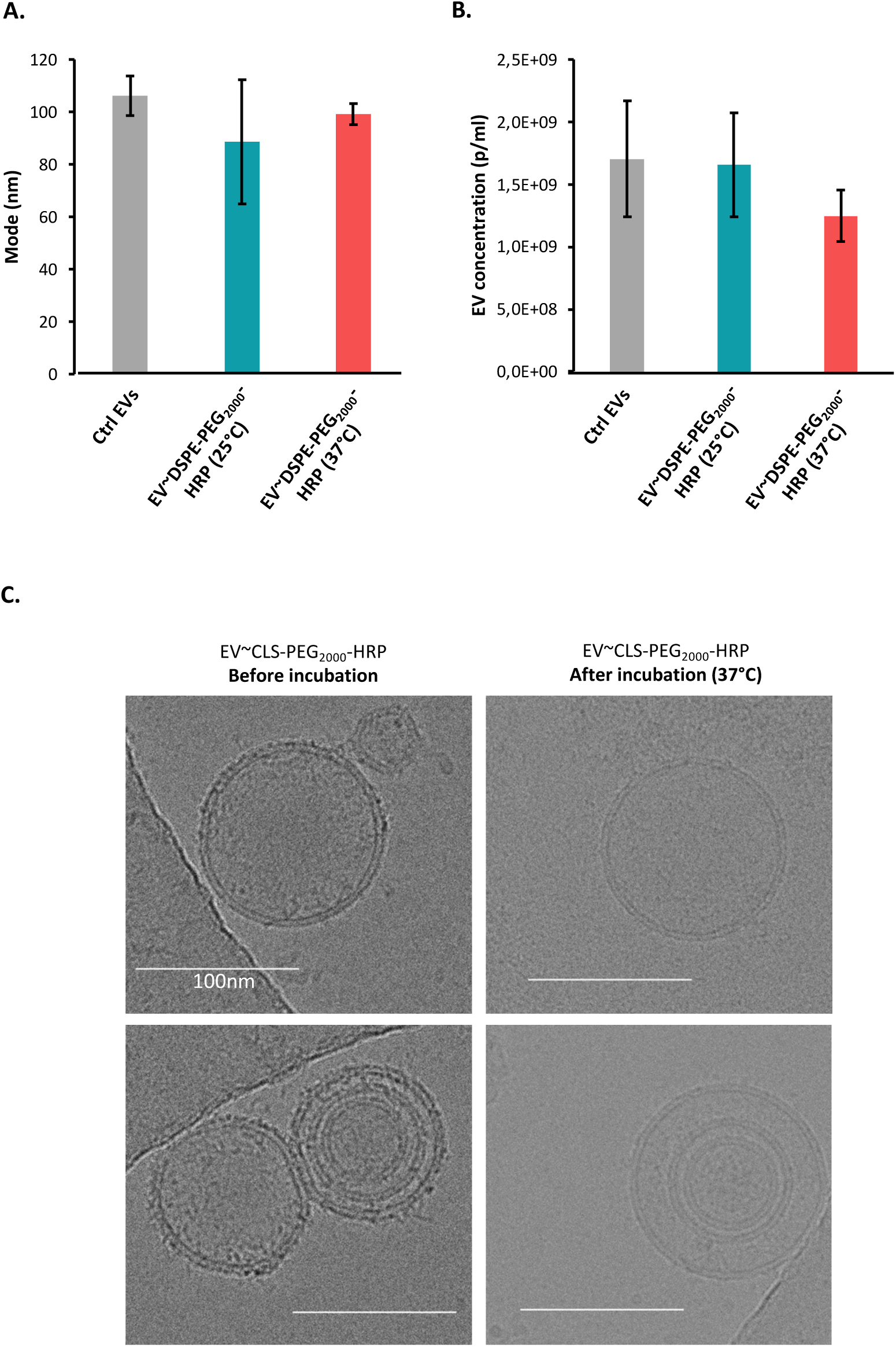
Influence of incubation temperature on EV size and concentration. EV mode **(A)** and concentration **(B)** after incubation (and subsequent SEC) with DSPE-PEG_2000_-HRP at 25°C or 37°C compared with unmodified ctrl EVs (NTA). n=3. **(C)** CryoTEM images of EV∼CLS-PEG_2000_-HRP before and after incubation at 37°C (no separation was processed). Bar scale is 100nm. For **(A)** and **(B)**, means and standard deviations are shown and the 4^th^ fraction of SEC has been fixed. p-values were calculated using a two-tailed Welch test. *n.s*>0.05.

**Supplementary Figure 4.**
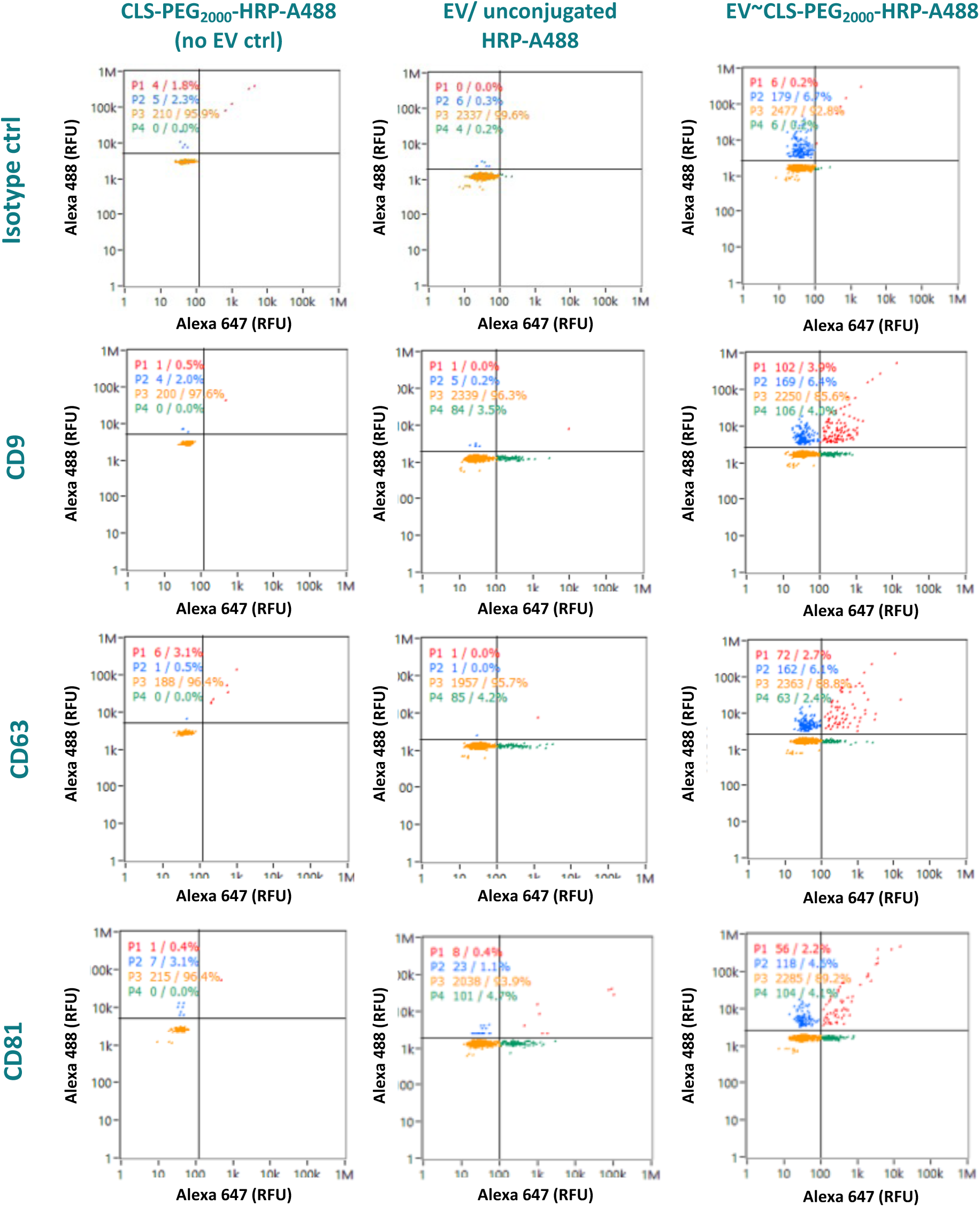
NanoFCM particle distribution of EV∼CLS-PEG_2000_-HRP-A488. Samples have been incubated with anti-CD9, anti-CD63 or anti-CD81 antibodies coupled with Alexa-Fluor 647. HRP was coupled with Alexa-Fluor 488.

**Supplementary Figure 5.**
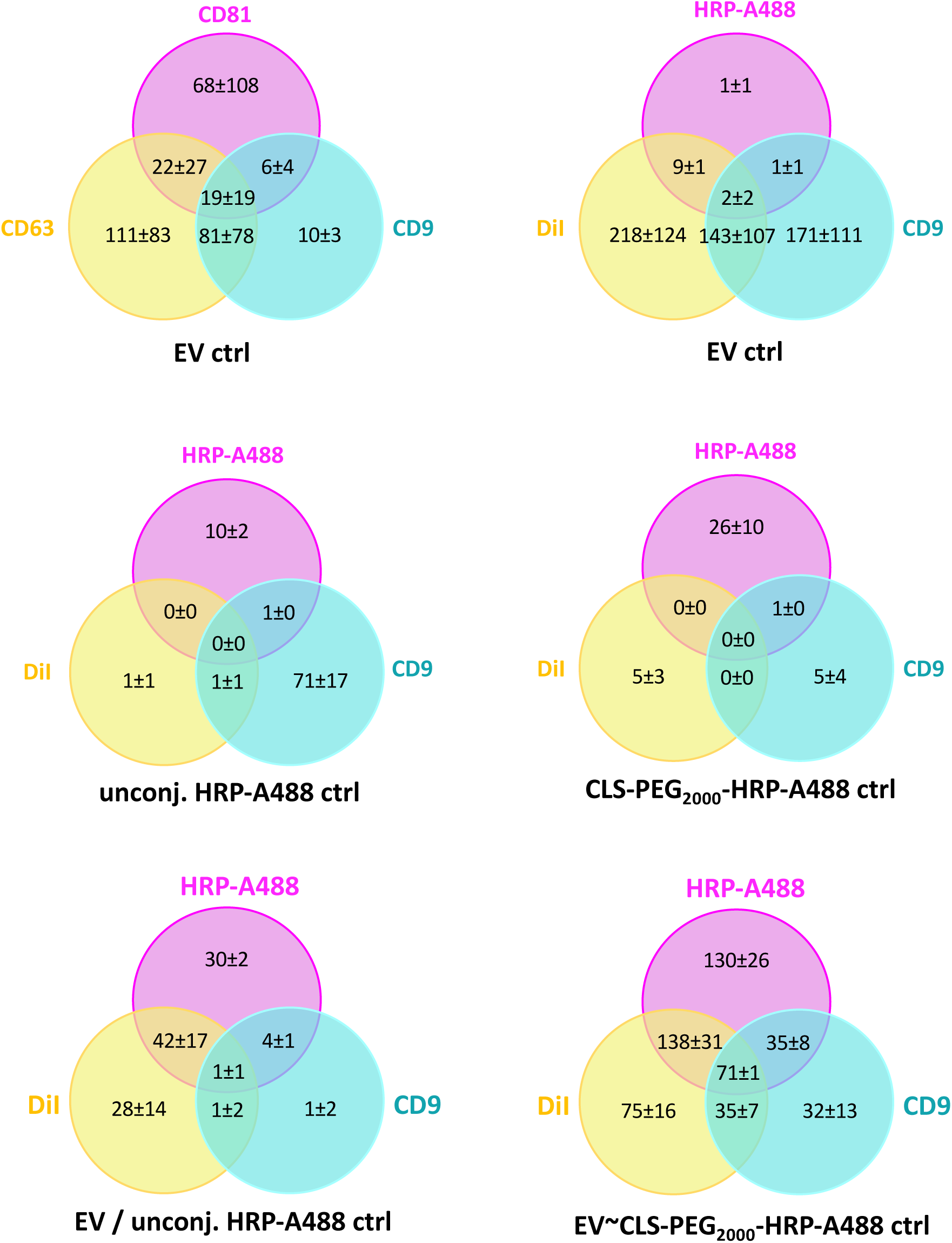
Super-resolution microscopy clusters distribution. Number of simple, double and triple positive counted clusters are shown (n=3). Samples were incubated with DiI dye and anti-CD9 antibody while HRP was labeled with Alexa-Fluor 488.

**Supplementary Figure 6.**
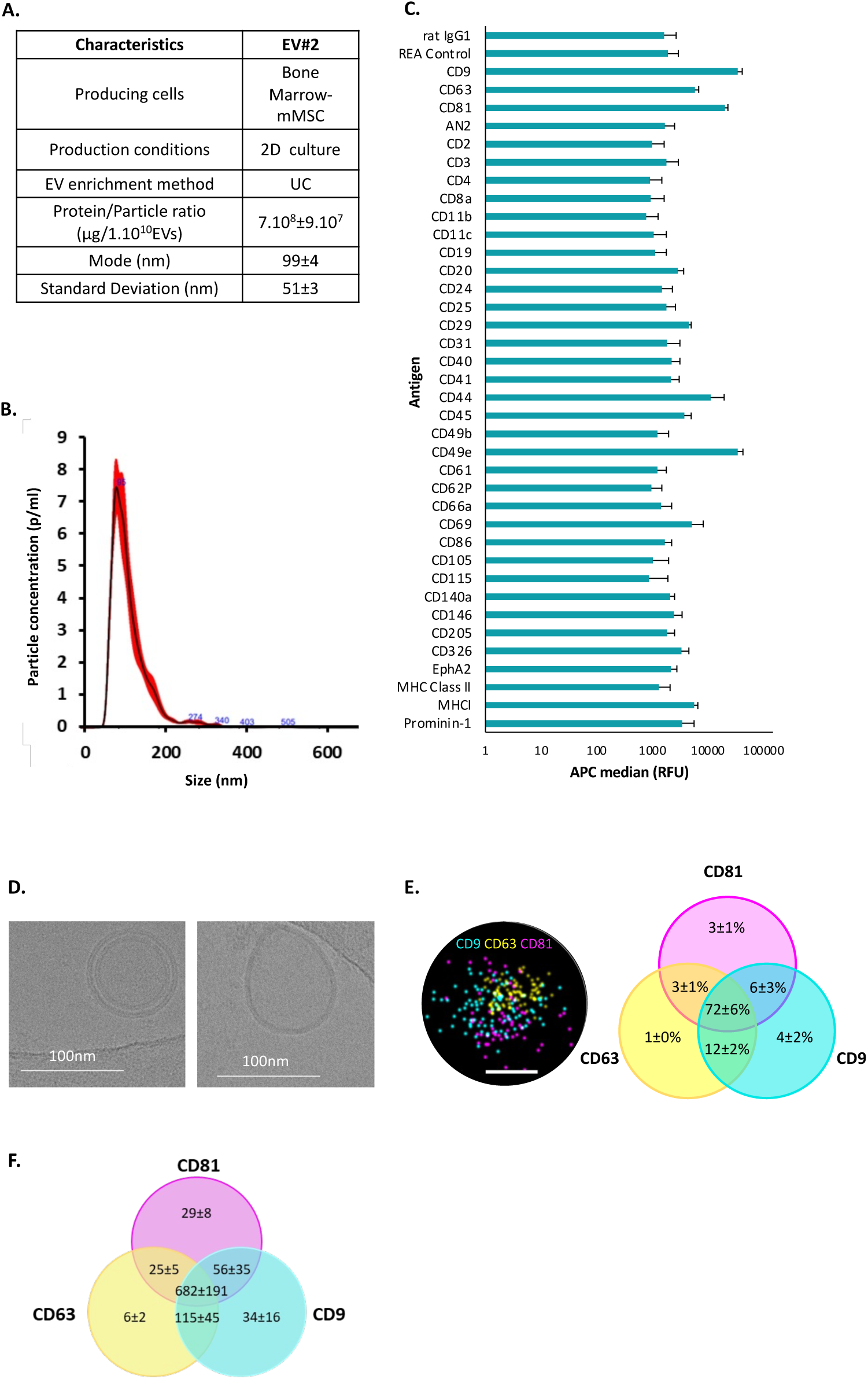
Characteristics of a second source of EVs (namely mMSC-EV#2) used to control the versatility of this approach. **(A)** Production and enrichment characteristics of mMSC-EV#2. Protein full content was measured using µBCA assay and BSA as reference (n=3). EV concentration, mode and SD were measured using NTA (NS300). **(B)** Size distribution profiles measured by NS300. **(C)** MACSPlex Assay signal as APC median after subtraction of buffer background signal (n=3). **(D)** CryoTEM images. **(E)** Proportion of single, double and triple positive clusters analyzed using STORM super-resolution microscopy. Samples were incubated with anti-CD9, -CD63 and -CD81 antibodies. Representative images of triple positive clusters are shown. Bar scale is 100nm (n=3). **(F)** ONI clusters distribution of the second source of EVs. Number of simple, double and triple positive counted clusters are shown (n=3). Samples were incubated with DiI dye and anti-CD9 antibody.

**Supplementary Figure 7.**
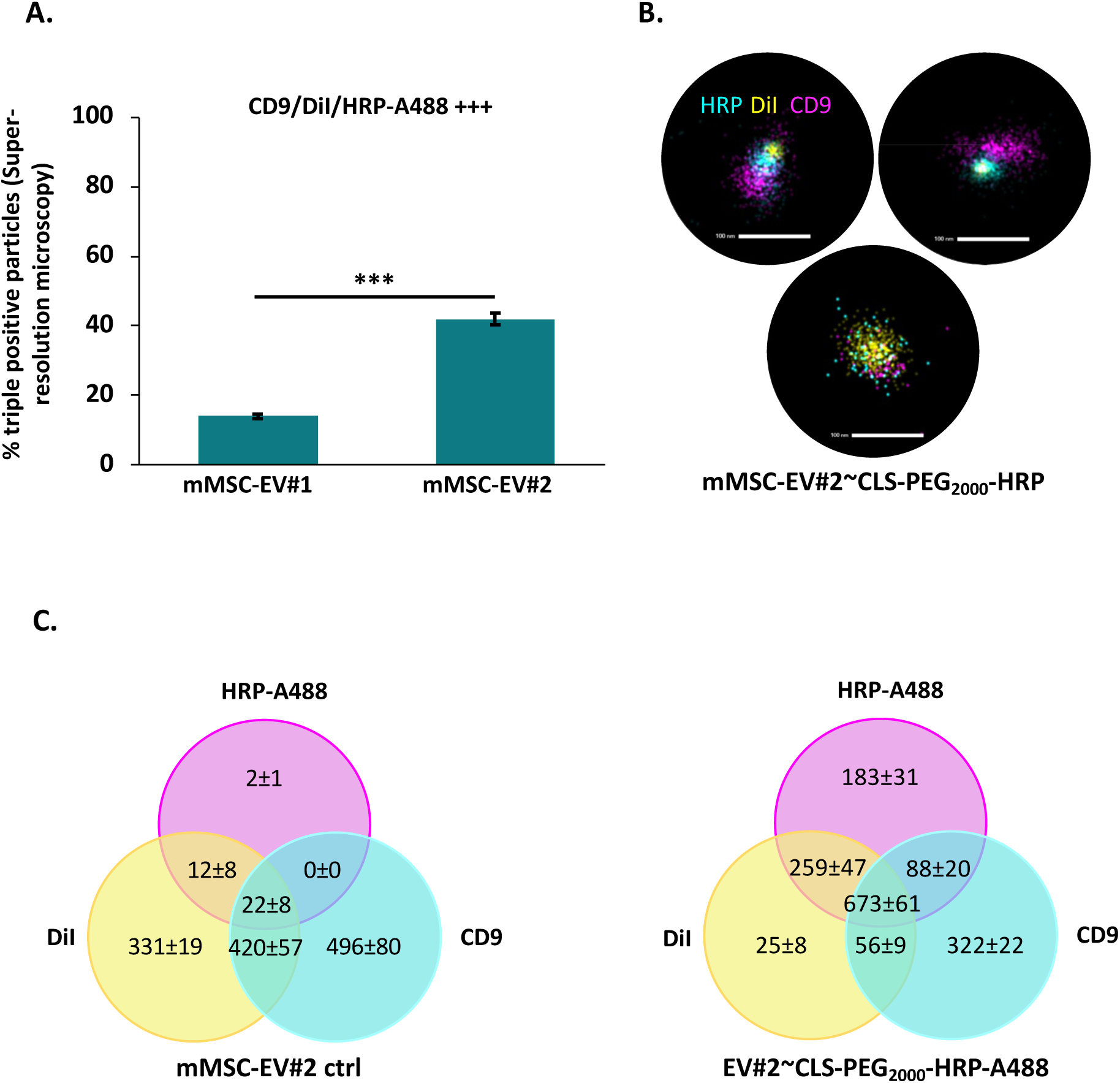
Investigation of the association of LA-HRP with a second source of EVs (mMSC-EV#2) using STORM. **(A)** Comparison of the proportion of triple positive clusters with the initial source of EVs used in this work (mMSC-EV#1) (n=3). **(B.)** Representative images of triple positive clusters of mMSC-EV#2 observed using super-resolution microscopy. **(C.)** ONI clusters distribution. p-values were calculated using a two-tailed Welch test. *n.s*>0.05, *<0.05, **<0,01, ***<0,001.

**Supplementary Figure 8.**
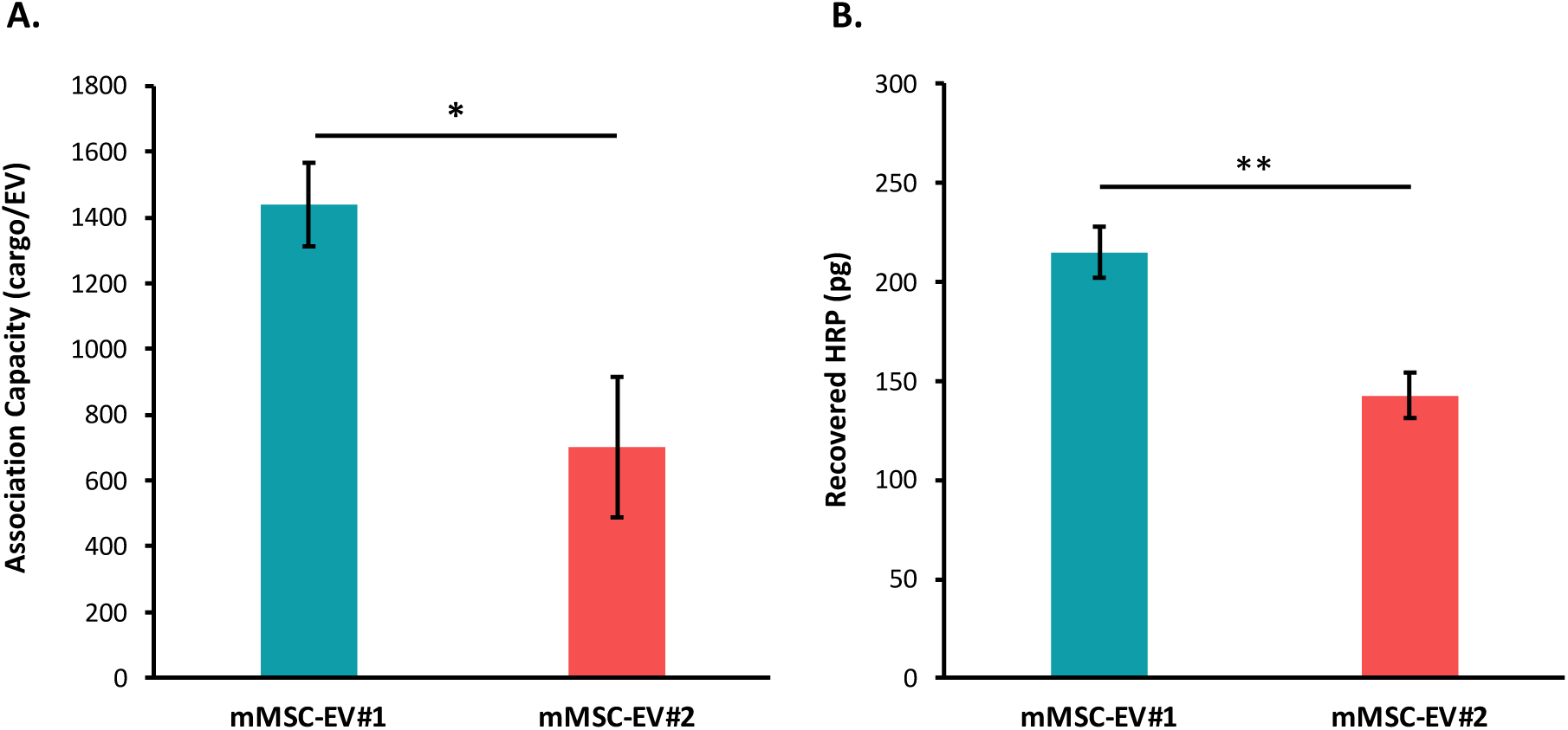
Comparison of the association and delivery capacity of mMSC-EV#1 (used in this study) and mMSC-EV#2 (additional EV product). **(A.)** Association capacity of CLS-PEG_2000_-HRP with mMSC-EV#1 or mMSC-EV#2. n=3. All EV SEC fractions (F3, F4 and F5) were considered (n=3). **(B.)** Quantification of absolute amount of internalized HRP by PANC-1 cells after incubation (4h, 37°C) of 10ng/well of EV∼CLS-PEG_2000_-HRP vectorized by mMSC-EV#1 or mMSC-EV#2 (n=3). p-values were calculated using a two-tailed Welch test. *n.s*>0.05, *<0.05, **<0.01.

## References

1. Tanaka, T. & Rabbitts, T. H. Interfering with protein-protein interactions: Potential for cancer therapy. Cell Cycle 7, 1569–1574 (2008).

2. Gulfidan, G., Turanli, B., Beklen, H., Sinha, R. & Arga, K. Y. Pan-cancer mapping of differential protein-protein interactions. Sci. Rep. 10, 3272 (2020).

3. Ivanov, A. A., Khuri, F. R. & Fu, H. Targeting protein–protein interactions as an anticancer strategy. Trends Pharmacol. Sci. 34, 393–400 (2013).

4. Maes, M., Loyter, A. & Friedler, A. Peptides that inhibit HIV-1 integrase by blocking its protein–protein interactions. FEBS J. 279, 2795–2809 (2012).

5. Naren, A. P., Quick, M. W., Collawn, J. F., Nelson, D. J. & Kirk, K. L. Syntaxin 1A inhibits CFTR chloride channels by means of domain-specific protein–protein interactions. Proc. Natl. Acad. Sci. 95, 10972–10977 (1998).

6. Poluri, K. M., Gulati, K., Tripathi, D. K. & Nagar, N. Protein–Protein Interactions in Immune Disorders and Inflammation. in Protein-Protein Interactions: Pathophysiological and Therapeutic Aspects: Volume II (eds. Poluri, K. M., Gulati, K., Tripathi, D. K. & Nagar, N.) 171–206 (Springer Nature, Singapore, 2023). doi:10.1007/978-981-99-2423-3_4.

7. Andrews, M. D. et al. Discovery of an Oral, Rule of 5 Compliant, Interleukin 17A Protein–Protein Interaction Modulator for the Potential Treatment of Psoriasis and Other Inflammatory Diseases. J. Med. Chem. 65, 8828–8842 (2022).

8. Gurevich, E. V. & Gurevich, V. V. Therapeutic Potential of Small Molecules and Engineered Proteins. in Arrestins - Pharmacology and Therapeutic Potential (ed. Gurevich, V. V.) 1–12 (Springer, Berlin, Heidelberg, 2014). doi:10.1007/978-3-642-41199-1_1.

9. Lu, R.-M. et al. Development of therapeutic antibodies for the treatment of diseases. J. Biomed. Sci. 27, 1 (2020).

10. Ebrahimi, S. B. & Samanta, D. Engineering protein-based therapeutics through structural and chemical design. Nat. Commun. 14, 2411 (2023).

11. Strohl, W. R. Current progress in innovative engineered antibodies. Protein Cell 9, 86–120 (2018).

12. Le Saux, S. et al. Interest of extracellular vesicles in regards to lipid nanoparticle based systems for intracellular protein delivery. Adv. Drug Deliv. Rev. 176, 113837 (2021).

13. Herrmann, I. K., Wood, M. J. A. & Fuhrmann, G. Extracellular vesicles as a next-generation drug delivery platform. Nat. Nanotechnol. 16, 748–759 (2021).

14. Wang, Z. et al. Exosomes decorated with a recombinant SARS-CoV-2 receptor-binding domain as an inhalable COVID-19 vaccine. Nat. Biomed. Eng. 6, 791–805 (2022).

15. Zhang, X. et al. Programmable Extracellular Vesicles for Macromolecule Delivery and Genome Modifications. Dev. Cell 55, 784–801.e9 (2020).

16. Lainšček, D. et al. Delivery of an Artificial Transcription Regulator dCas9-VPR by Extracellular Vesicles for Therapeutic Gene Activation. ACS Synth. Biol. 7, 2715–2725 (2018).

17. Ilahibaks, N. F. et al. TOP-EVs: Technology of Protein delivery through Extracellular Vesicles is a versatile platform for intracellular protein delivery. J. Controlled Release 355, 579–592 (2023).

18. Heath, N. et al. Endosomal escape enhancing compounds facilitate functional delivery of extracellular vesicle cargo. Nanomed. 14, 2799–2814 (2019).

19. Mizrak, A. et al. Genetically Engineered Microvesicles Carrying Suicide mRNA/Protein Inhibit Schwannoma Tumor Growth. Mol. Ther. 21, 101–108 (2013).

20. Haney, M. J. et al. TPP1 Delivery to Lysosomes with Extracellular Vesicles and their Enhanced Brain Distribution in the Animal Model of Batten Disease. Adv. Healthc. Mater. 8, e1801271 (2019).

21. Silva, A. M. et al. Quantification of protein cargo loading into engineered extracellular vesicles at single-vesicle and single-molecule resolution. J. Extracell. Vesicles 10, e12130 (2021).

22. Yim, N. et al. Exosome engineering for efficient intracellular delivery of soluble proteins using optically reversible protein–protein interaction module. Nat. Commun. 7, 12277 (2016).

23. Wang, Q. et al. ARMMs as a versatile platform for intracellular delivery of macromolecules. Nat. Commun. 9, 960 (2018).

24. Lamichhane, T. N., Raiker, R. S. & Jay, S. M. Exogenous DNA Loading into Extracellular Vesicles via Electroporation is Size-Dependent and Enables Limited Gene Delivery. Mol. Pharm. 12, 3650–3657 (2015).

25. Corso, G. et al. Systematic characterization of extracellular vesicle sorting domains and quantification at the single molecule – single vesicle level by fluorescence correlation spectroscopy and single particle imaging. J. Extracell. Vesicles 8, 1663043 (2019).

26. Ivanova, A. et al. Creating Designer Engineered Extracellular Vesicles for Diverse Ligand Display, Target Recognition, and Controlled Protein Loading and Delivery. Adv. Sci. 10, 2304389 (2023).

27. Rankin-Turner, S. et al. A call for the standardised reporting of factors affecting the exogenous loading of extracellular vesicles with therapeutic cargos. Adv. Drug Deliv. Rev. 173, 479–491 (2021).

28. Jing, B. et al. Extracellular vesicles-based pre-targeting strategy enables multi-modal imaging of orthotopic colon cancer and image-guided surgery. J. Nanobiotechnology 19, 151 (2021).

29. Zheng, W. et al. Surface display of functional moieties on extracellular vesicles using lipid anchors. J. Controlled Release 357, 630–640 (2023).

30. Choi, E. S., Song, J., Kang, Y. Y. & Mok, H. Mannose-Modified Serum Exosomes for the Elevated Uptake to Murine Dendritic Cells and Lymphatic Accumulation. Macromol. Biosci. 19, 1900042 (2019).

31. Pitchaimani, A. et al. Compartmentalized drug localization studies in extracellular vesicles for anticancer therapy. Nanoscale Adv. 5, 6830–6836 (2023).

32. Didiot, M.-C. et al. Exosome-mediated Delivery of Hydrophobically Modified siRNA for Huntingtin mRNA Silencing. Mol. Ther. 24, 1836–1847 (2016).

33. Haraszti, R. A. et al. Optimized Cholesterol-siRNA Chemistry Improves Productive Loading onto Extracellular Vesicles. Mol. Ther. 26, 1973–1982 (2018).

34. Stremersch, S. et al. Comparing exosome-like vesicles with liposomes for the functional cellular delivery of small RNAs. J. Controlled Release 232, 51–61 (2016).

35. Zhang, H. et al. Exosome-mediated targeted delivery of miR-210 for angiogenic therapy after cerebral ischemia in mice. J. Nanobiotechnology 17, 29 (2019).

36. Gong, C. et al. Functional exosome-mediated co-delivery of doxorubicin and hydrophobically modified microRNA 159 for triple-negative breast cancer therapy. J. Nanobiotechnology 17, 93 (2019).

37. Biscans, A. et al. Hydrophobicity of Lipid-Conjugated siRNAs Predicts Productive Loading to Small Extracellular Vesicles. Mol. Ther. 26, 1520–1528 (2018).

38. Tréton, G. et al. Quantitative and functional characterisation of extracellular vesicles after passive loading with hydrophobic or cholesterol-tagged small molecules. J. Controlled Release 361, 694–716 (2023).

39. Hagey, D. W. et al. The cellular response to extracellular vesicles is dependent on their cell source and dose. Sci. Adv. 9, eadh1168 (2023).

40. Rennke, H. G. & Venkatachalam, M. A. Chemical modification of horseradish peroxidase. Preparation and characterization of tracer enzymes with different isoelectric points. J. Histochem. Cytochem. Off. J. Histochem. Soc. 27, 1352–1353 (1979).

41. Reinhardt, K. Über die dichteste gitterf örmige lagerung kongruenter bereiche in der ebene und eine besondere art konvexer kurven. Abh. Aus Dem Math. Semin. Univ. Hambg. 10, 216–230 (1934).

42. Fejes Tóth, L., Fejes Tóth, G. & Kuperberg, W. Lagerungen: Arrangements in the Plane, on the Sphere, and in Space. vol. 360 (Springer International Publishing, Cham, 2023).

43. Chen, C., et al. Active cargo loading into extracellular vesicles: Highlights the heterogeneous encapsulation behaviour. J. Extracell. Vesicles 10, e12163 (2021).

44. Mitrut, R. E., et al. HaloTag display enables quantitative single-particle characterisation and functionalisation of engineered extracellular vesicles. J. Extracell. Vesicles 13, e12469 (2024).

45. van Niel, G., D’Angelo, G. & Raposo, G. Shedding light on the cell biology of extracellular vesicles. Nat. Rev. Mol. Cell Biol. 19, 213–228 (2018).

46. Mizenko, R. R. et al. Tetraspanins are unevenly distributed across single extracellular vesicles and bias sensitivity to multiplexed cancer biomarkers. J. Nanobiotechnology 19, 250 (2021).

47. Alwarawrah, M., Dai, J. & Huang, J. A Molecular View of the Cholesterol Condensing Effect in DOPC Lipid Bilayers. J. Phys. Chem. B 114, 7516–7523 (2010).

48. Skotland, T., Sagini, K., Sandvig, K. & Llorente, A. An emerging focus on lipids in extracellular vesicles. Adv. Drug Deliv. Rev. 159, 308–321 (2020).

49. Le Saux, S. et al. Post-production modifications of murine mesenchymal stem cell (mMSC) derived extracellular vesicles (EVs) and impact on their cellular interaction. Biomaterials 231, 119675 (2020).

50. Tulkens, J., De Wever, O. & Hendrix, A. Analyzing bacterial extracellular vesicles in human body fluids by orthogonal biophysical separation and biochemical characterization. Nat. Protoc. 15, 40–67 (2020).

